# Implicit Adaptation is Fast, Robust and Independent from Explicit Adaptation

**DOI:** 10.1101/2024.04.10.588930

**Authors:** Sebastian D’Amario, Jennifer E. Ruttle, Bernard Marius ’t Hart, Denise Y. P. Henriques

## Abstract

During classical visuomotor adaptation, the implicit process is believed to emerge rather slowly; however, recent evidence has found this may not be true. Here, we further quantify the time-course of implicit learning in response to diverse feedback types, rotation magnitudes, feedback timing delays, and the role of continuous aiming on implicit learning. We find that implicit learning unfolds at a higher rate than conventionally expected in all feedback conditions. Increasing rotation size not only raises asymptotes, but also generally heightens explicit awareness, with no discernible difference in implicit rates. Cursor-jump and terminal feedback, with or without delays, predominantly enhance explicit adaptation while slightly diminishing the extent or the speed of implicit adaptation. In a continuous aiming reports condition, there is no discernible impact on implicit adaptation, and implicit and explicit adaptation progress at indistinguishable speeds. Finally, investigating the assumed negative correlation as an indicator of additivity of implicit and explicit processes, we observe weak associations at best across conditions. Our observation of implicit learning early in training in all tested conditions signifies how fast and robust our innate adaptation system is.

## Introduction

People constantly adapt their movements to changing circumstances, and this adaptation is driven by a combination of implicit and explicit processes. Implicit motor learning offers the advantage of preserving cognitive resources, thereby boosting performance efficiency. In contrast, employing explicit strategies demands more effort and may prove less efficient over the long term, even though it may manifest more quickly than implicit contributions to the learning process. People rely more on implicit processes when carrying out well-learned motor tasks, but it is harder to quantify these unconscious contributions to our performance, or even identify when they emerge and under what conditions. This study aims to investigate the time-course of implicit contributions during classical visuomotor adaptation and explore its sensitivity to different kinds of visual feedback.

Visuomotor adaptation is classically studied by having participants reach to targets with a misaligned hand-cursor that misrepresents their unseen hand. People rapidly adjust their reach direction in response to the deviated cursor motion. It is assumed that the initial compensation, achieved by directing the unseen hand elsewhere to move the cursor to the target, is driven by explicit strategy. Implicit contributions to adaptation are thought to emerge later and gradually replace the cognitive strategy as the movements become more automatic. Implicit adaptation is traditionally measured through reach aftereffects, which refer to the residual deviations in subsequent reaching movements even after the feedback is removed or returned to normal.

Recently, some studies [1,2] have used “clamped error feedback” to assess implicit adaptation. In such paradigms, the cursor will always move in a straight line from the start position in a direction that misses the target by some predetermined amount. The cursor’s distance from the home position typically matches the hand’s distance from the home position, such that participants do feel some measure of control over the cursor. This situation where reaches always result in the same error is combined with instructions to participants to ignore this feedback and to keep moving the hand to the target as opposed to moving the cursor to the target. That is, participants are told to disregard and not learn from the only visible feedback they receive on their performance. Despite participants’ best efforts to suppress any and all learning, their reaches do slowly start to deviate from the target. Eventually they reach levels of adaptation comparable to implicit reach aftereffects recorded after a prolonged learning task.

While the data from these types of paradigms are impressive reminders of the power of our implicit motor adaptation system, the participants’ efforts to suppress learning likely reduces the speed of implicit adaptation, perhaps by as much as an order of magnitude. Here, we assess the speed of implicit adaptation without any suppression.

Studies using reach aftereffects and error-clamp paradigms indicate that implicit adaptation tends to saturate at around 10-20° independent of the rotation size [1–3]. Because these implicit changes have been assumed to emerge much later, reach aftereffects are usually not measured until at least 60 or more perturbed training trials. However, recent research from our lab suggests that substantial implicit changes in hand movement can emerge rapidly within the first few trials of training with a visuomotor rotation [4]. This paper will focus on testing implicit learning rates during visuomotor adaptation using our method of interleaving visual cursor feedback trials with no-cursor feedback trials [4].

While implicit and explicit processes both contribute to adaptation, the way they interact remains unclear [5]. Hence, in this study, we examine implicit learning rates and asymptotes across various paradigms that are thought to increase, explicit adaptation. This allows mapping out how implicit adaptation responds to changes in explicit adaptation. We use the rate of change of an exponential learning function fitted to no-cursor reach deviations to assess the rate of implicit learning. By fitting this function to all no-cursor trials before any other change in the task occurs, we maximally reduce the effect noise on our estimates of learning rate.

First we test whether varying the cursor rotation size changes the rate of implicit adaptation. Previous research has indicated that the extent of implicit learning does not necessarily scale linearly with the magnitude of the perturbation, but seems to be capped at roughly 15° [1,6,7]. However, it remains unclear whether the time course by which aftereffects emerge is consistent across different rotation sizes as well.

The second study investigates the impact of feedback types. Specific kinds of visual feedback are known to affect explicit strategy use during visuomotor adaptation. We will employ two manipulations: terminal feedback and cursor jump. Terminal feedback offers limited visual feedback, providing cursor information only at the reach’s endpoint (Fig 4). Cursor jump alters the cursor’s path mid-movement, creating a salient error signal that encourages strategy use (Fig 4). Previous research has produced mixed findings on the impact of terminal feedback on adaptation. Some studies [8–12] suggest that terminal training may slow down adaptation and reduce aftereffects, while others [13,14] find no significant differences. However, all of these studies typically assess aftereffects after 60 to 100 trials, making it unclear how the rate of these implicit changes develops over time. Cursor jump raises awareness of external perturbation [15], implying more reliance on explicit strategy. In this study by Gastrock et al, smaller implicit reach aftereffects were found following 90 trials of training. Thus, developing an explicit strategy during cursor-jump training could lead to a reduction in implicit-driven changes, and it is possible that it could also delay the onset of reach aftereffects. That is, both terminal feedback and cursor jump feedback seem to decrease the extent of implicit adaptation, but the effect of these types of feedback on the speed of implicit adaptation is unknown. We test that here.

Following this, we sought to observe the temporal progression of implicit processes when employing a paradigm designed to minimize their influence. Studies have found that the implicit component is reduced, although not entirely eliminated, when the terminal feedback of the cursor is delayed by varying durations: 5 and 1 seconds [9], 1.5 seconds [14], and 1.1-1.3 seconds [16]. In our Feedback-delay study, we aimed to investigate whether aftereffects decrease with a 1.2-second delay in feedback during training and whether there is a hindrance in their onset.

In all of the aforementioned conditions, we not only assessed reach aftereffects with a high level of temporal precision but also intermittently gathered aiming trials during the latter stages of training to evaluate explicit contributions to adaptation [6,12,17–19]. For our final study, we introduced a condition in which we measured both reach aftereffects and aiming at the same high rate. This allowed us to compare the rate of changes across both, without relying on any assumptions about the relationship between the two kinds of processes, e.g., without assuming that implicit and explicit contributions are necessarily additive.

Collecting frequent aiming responses and implicit measures allows us to determine the extent to which these two processes are interconnected in the learning process [5]. While implicit adaptation has been explored through clamp trials [2,20,21], our approach involves measuring the type of implicit motor changes that contribute to the types of motor adaptation we routinely experience when interacting with our dynamically changing environments. By understanding the time course of these more natural implicit changes during motor learning, we can gain valuable insights that will help us enhance training and adaptation for various real-world scenarios.

## Methods

### Participants

We used data from 378 volunteer participants. These all had normal or corrected-to-normal vision (mean age = 21, females = 239) from the Undergraduate Research Participation Pool (URPP), and the Kinesiology Undergraduate Research Experience (KURE), who all provided prior, written, informed consent. The procedures used in this study were approved by York University’s Human Participant Review Committee and all experiments were performed in accordance with institutional and international guidelines.

Participants were randomly assigned to 10 experimental groups, first to the groups in the feedback type experiment (n=126), then to the delayed feedback group and its control (n=67) and the rotation size experiment groups (n=176), and finally to the continuous aiming group (n=37). We can only assess the speed of any adaptation process if there is some amount of adaptation. This is why we only used data from participants whose reach deviations in the last 20 trials of the rotated phase are on average countering at least 50% of the rotation in their condition (i.e. we do not select participants based on no-cursor reach deviations, the main measure of interest, nor based on aiming responses). The participants listed at the start of the methods and used in the analyses did meet our criterion, however we had to remove 15 participants in the *Feedback Type* experiment, 18 participants from the *Feedback Delay* experiment, 43 participants from the *Rotation Size* experiment, and 4 participants from the *Continuous Aiming* group. The data from *all* participants is available on an OSF repository (https://osf.io/ajwyr/).

### Experimental Set-up

#### Apparatus

Participants sat on a height-adjustable chair facing a digitizing tablet (Wacom Intuos3, 12” x 12” surface, sampled on every frame refresh) and screen (Fig 1A). The tablet was positioned at waist level so hand movements could be made along a horizontal plane (See Fig 1A for detail). On top of the tablet there was a stencil with a circular portion cut-out measuring 20 cm in diameter (further details found on OSF: https://osf.io/7pzrb/). Visual feedback was shown on a computer screen located approximately 60 cm from the tablet workspace (22” monitor, 1680×1050 pixels, 60 fps). A wooden shield was placed above the tablet work surface to obstruct participants’ view of their arm movements. Participants used a digital stylus to move the cursor (0.7 cm in diameter) onto the target displayed on a vertical screen (Fig 1A). The trial began when the cursor was moved to the home position. Participants had to move the stylus 8.8 cm to reach the target, with a margin up to the edge of the stencil of 1.2 cm. The stencil, positioned atop the digital tablet, effectively restricted the radial movement of reaches toward the target. It achieved this by physically impeding any outward movement beyond this limit.

**Fig 1.**
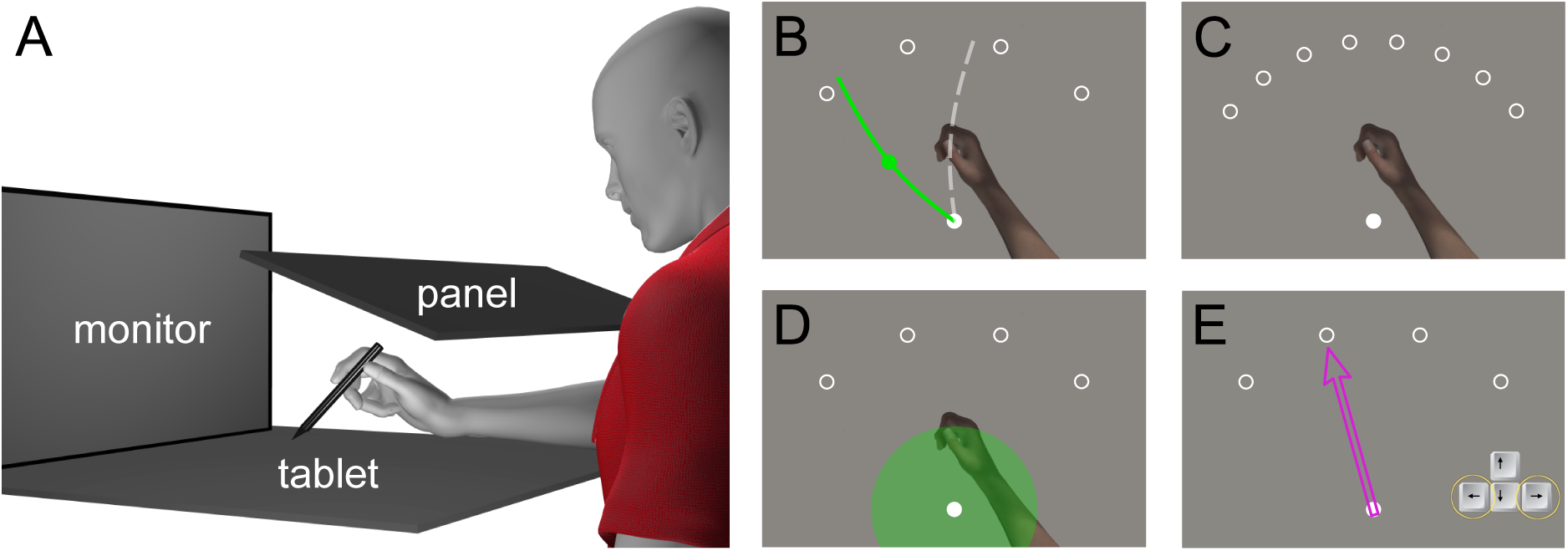
**A.** Setup for all experiments. The stylus slides over a digitizing tablet while pen movements correspond to cursor movements on the connected upright screen. Hand view is blocked by a wooden panel. **B.** Reach training trial in the rotated phase of the experiment where the cursor position corresponded to the stylus position, rotated at the set perturbation depending on which condition the participant was assigned to. **C.** No-cursor trials would involve making a reach with no cursor feedback out to one of 8 targets. **D.** When performing the no-cursor trials, participants would see a green disk, which would help guide their movements out to the target as it informed them of their distance from the home position. **E.** In aiming trials, participants used left and right arrow keys to move the arrow to point in a direction they were moving their hand to hit the target.

### Trial Types

For all 10 conditions, participants experienced a very similar main trial structure within the experiment. This involved alternating cursor trials (with further feedback details given in each experiment) with no-cursor trials (Fig 1C-D) where participants were told to directly move their unseen hand to the target. Later on in the experiment’s rotated phase, we included eight aiming trials. Only in the *Continuous Aiming* experiment did we have all three of these trials in succession of one another (aiming trial - cursor trial - no-cursor trial) for the entire experiment. Each of these components will be discussed below. We colour-coded the cursor and target to provide participants with extra feedback about the trial type as well as their performance. In both aligned and rotated phases (as well as washout), the cursor was white during reaching trials, and in the no-cursor trials the target was green. In the reach/training trials, the target was green when the cursor was aligned, and purple when the cursor was rotated. We also had the target change colour after the outward reach was completed to signify to the participant if they performed the trial according to our criteria outlined below. The target would turn blue when they met the criteria, and orange if they did not.

### Cursor Trials

Cursor, or Reach-training trials involve participants making out-and-back reaching movements to a target with the cursor. All conditions used one of four forward targets on each trial (0.7 cm in diameter) located at 45°, 75°, 105°, and 135° as shown in Figure 1B. Participants reached to the target, and upon completion, the target would vanish, and participants would receive feedback about movement accuracy, and then move their cursor back to the home position.

### No-Cursor Trials

These trials worked very similarly to the *Cursor Trials* with two notable differences. Primarily, participants were unable to see their hand position during the outward reach. Instead of a cursor, they used a green disc (shaded filled circle) which was centered on the home position and increased in size the farther away they moved from the home position during their outward reach. Subsequently, participants returned their hand to the home position without a cursor, with the assistance of the green disc that indicated the remaining distance to the home position.

When the tip of the stylus was within 3.5 cm of the home position, the cursor became visible again to ensure a precise return to home. Second, the target alternated between eight different locations, such that the current no-cursor target was located ± 7.5° degrees from the previous reach-training target. These locations were 37.5°, 52.5°, 67.5°, 82.5°, 97.5°, 112.5°, 127.5° and 142.5° (as shown in Fig 1C). This was meant to prevent participants from simply repeating the reaching movement from the previous training trial.

### Aiming Trials

Aiming trials (also called re-aiming), shown in Fig 1E far right, are used to measure the explicit component of adaptation. Participants adjusted the arrow’s direction using the left- and right arrow keys to indicate the direction they planned to move their hand so that the cursor would hit the target. Once they got to the desired position, they would press the spacebar, and would be able to continue with the next reach-training trial. Following previous studies, we used aiming trials to assess the extent that adaptation reflects a cognitive strategy in all conditions. Crucially, the participants were not told about the arrow trials before they adapted so as to prevent the concepts present in the instructions to aiming trials (“in which direction would you move your hand in order to make the cursor hit the target”), as well as performing the aiming trials themselves from increasing strategy-based adaptation. We withheld the instruction until it was needed, to ensure that our measure of the natural time-course of implicit adaptation was not affected by the instructions about the aiming responses. Participants were given on-screen instructions before the aiming trials appeared. As expected, this led to some de-adaptation right after the first few aiming trials. While participants quickly recovered, this is why we included them later in the experiment so as not to affect the fitted initial change calculation. Of course, in the continuous aiming condition, the instructions were given in the familiarization phase and never re-appeared.

### General Procedure

After providing informed consent and demographic information, all participants watched a basic instruction video in an effort to standardize the instructions received (these are all on OSF: https://osf.io/ajwyr/). They were allowed an opportunity to ask questions if something in the video was unclear or they could re-watch the video. The experiment consisted of three distinct phases, a practice phase for familiarization with the task, an aligned phase where baseline performance was established, and a rotated phase when the perturbation was introduced. For the first experiment, we also included a fourth washout phase to measure de-adaptation of the rotation.

Participants began by completing a practice phase of the experiment, consisting of 16 reach-training trials, and the interleaved 16 no-cursor trials. If the condition involved terminal feedback during reach-training trials or continuous aiming, this was introduced in the second half of the practice phase. During the practice phase, participants were given feedback on their reaching movements. They would see “too slow!” if they did not complete the reach within 1500 ms, and “missed target!” if the cursor missed the target by 15° or more (10° in the condition with a 15° rotation). Throughout the entire experiment, the target was initially green or purple, but would turn blue if participants performed the trial correctly, and turn orange if it was done incorrectly, based on the criteria above. This colour feedback about performance was provided both to guide performance and to keep participants motivated.

Before the real task started, there was a break that allowed for any remaining questions. The experiment then started with an aligned phase of 20 reach-training trials and the interleaved 20 no-cursor trials. In both the practice phase and aligned phase, the direction of the hand-cursor motion was aligned with the unseen hand. After participants completed the 16th aligned pair of trials, a warning screen with instructions was shown telling them that in eight trials the original green target was going to turn purple and that they were going to have to move a bit differently for the cursor to hit the target. Most importantly, they were told to not slow down, but to keep making reaches at the established pace, and that they should keep making straight reaching movements. This was also mentioned to them in the instruction video prior to beginning the experiment. This colour change to purple marked the beginning of the rotated phase. In this, they would complete 100 pairs of trials with alternating reach-training trials and no-cursor trials. In the reach-training trials presented in this phase the cursor was shown at a location that represented the stylus position, rotated about the starting position. The exact amount of rotation differed across groups as described below, but was 45° in most groups. The no-cursor trials were the same as before, and participants were told to reach directly to the target. Before beginning the 57th pair of rotated trials, participants performed the first aiming trial, and did so 7 more times, once after every 4 pairs of trials. All experiments were monitored by a Research Assistant to confirm that participants followed the instructions provided.

### Data Analysis

Our experiment has two different trial types with similar analysis methods used in each. In both cursor and no-cursor trials, participants completed an out-and-back reach for which we calculated the deviation of the outward reach from a straight reach to the target. For both, we took the first sample further than 1.8 cm from the home position, and calculated the angle between a line through the home position and this sample and a line through the home position and the target. To quantify explicit awareness, we took the direction of the arrow relative to the direction of the target. In all aiming trials, the arrow started out 15° CCW relative to the target. Since in all rotations people moved their unseen hand in a direction CW to compensate for CCW rotation, participants *without* a strategy would need to move the arrow approximately to the target, and participants *with* a strategy would need to move it further CW. We rejected arrow aiming directions that were unreasonable, specifically those where the arrow did not move more than 5° from its original 15° CCW direction or moved in the wrong direction relative to where their unseen hand should have moved to compensate for the visuomotor rotation, as well as outliers larger than 120°. We did so because in these trials it is likely the participant either did not move the arrow due to a misunderstanding or erroneously ended the aiming trial.

For each dependent measure, reach deviations for reach-training trials and no-cursor trials and aiming deviations for aiming trials, a 0 indicates no adaptation and a value equal to the rotation would indicate full adaptation.

### Analyses

We used the aligned phase, excluding the initial four trials, to determine each participant’s baseline biases by calculating the median reach deviation separately for trials with and without the cursor. These biases were then subtracted from all corresponding trials in the aligned and rotated phases, and the washout phase of the rotation size experiment. Prior to removing participants who did not demonstrate learning, we performed outlier removal and baselining.

Outliers for reaches (both with and without cursors) were identified as deviations more than 15 degrees in the incorrect direction or exceeding the rotation plus 15 degrees in the correct direction. For aiming responses, outliers were those more than 10 degrees in the wrong direction or more than twice the rotation value (multiplied by −1). Unlike reach trials, aiming responses were not baselined. For reaches, the last 16 trials of the baseline period were used to calculate the median reach, excluding any trials identified as outliers. This process ensured the exclusion of non-learners was based on reliable and standardized performance measures across all trial types.

We wanted to investigate the rate of change in reach training trials, implicit no-cursor test trials, and of aiming responses. However, to determine a rate of change we first need to validate that participants did change their performance over the course of training with the rotation. This is why in all groups we checked if participants were compensating for at least 50% of the perturbation on average in the last 20 trials of the rotated phase. After this criteria was met, we retained data from 378 out of 458 participants.

### Exponential Learning Function for Rate of Change

To rigorously quantify the time-course of the implicit process we used an exponential learning function which used error decay with an asymptote to identify a rate of change for each trial type in each feedback condition. We used the same equation, shown below, as used previously [4]. The value of the process on the current trial (P_t_) is the process’ value on the previous trial (P_t-1_) plus the product of the rate of change (L) multiplied by the error on the previous trial, which is the difference between the asymptote (A) and the process’ value on the previous trial (P_t-1_).

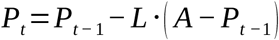

The parameter L was constrained to the range [0,1], and the parameter A to [-1,2]*max(data). This model was fit to the rotated reach data and reach aftereffect data for all groups. In order not to overestimate the speed at which aftereffects arise, a zero was prepended to this time course. This accounts for the fact that responses in these trials already changed through the previous training trial. Each parameter was bootstrapped (5 k resamples per fit) across participants to get a 95% confidence interval which can then be compared, and values can be found in Tables 1-4. Whenever plotted, the line in the middle of the shaded area represents a central value—the median, which corresponds to the 50% value of the bootstraps.

**Table 1.** Descriptives of adaptation for each group in the rotation size experiment. Rates of change (RofC) and asymptotes are shown for training and no-cursor reaches in the rotated phase. Aiming extent is shown as the average across the eight aiming trials in the second half of the rotated phase. For the washout phase, only the start level of functions fitted to training and no-cursor reaches are shown (not the retention rates, which are all between 87% and 95% and not different between conditions). The entries for RofC, asymptote and start level all first list (what could be considered) the group average using each participant once, then in parentheses the 95% confidence interval of the mean, based on 5k bootstraps across participants. For RofC we also list the absolute value after one rotated trial as predicted by the function fitted to all data.

|  |  |  | 15° (N=33) | 30° (N=54) | 45° (N=28) | 60° (N=61) |
| --- | --- | --- | --- | --- | --- | --- |
| Rotated phase | Reach training | RofC | 19.9%<br>(14.8%-27.2%)<br>2.0° | 19.8%<br>(15.2%-25.6%)<br>3.7° | 18.3%<br>(12.5%-28.3%)<br>5.8° | 14.5%<br>(10.9%-20.0%)<br>6.1° |
|  |  | asymptote | 10.0°<br>(9.0°-11.0°) | 18.6°<br>(17.1°-20.2°) | 31.9°<br>(28.3°-35.4°) | 41.8°<br>(39.1°-44.5°) |
|  | No-cursors | RofC | 22.9%<br>(16.7%-32.6%)<br>2.4° | 18.5%<br>(14.6%-23.2%)<br>3.6° | 11.0%<br>(7.9%-16.5%)<br>2.7° | 8.8%<br>(7.0%-11.1%)<br>2.4° |
|  |  | asymptote | 10.3°<br>(9.2°-11.5°) | 19.6°<br>(18.2°-21.0°) | 24.8°<br>(21.4°-28.5°) | 26.9°<br>(24.3°-29.7°) |
|  | Re-aiming | extent | 0.9°<br>(-0.2°-2.2°) | 3.7°<br>(2.4°-5.1°) | 13.7°<br>(9.8°-18.3°) | 21.3°<br>(17.4°-25.3°) |
| Washout phase | Reach training | start level | 9.8°<br>(8.4°-11.3°) | 18.8°<br>(17.0°-20.7°) | 20.0°<br>(16.3°-23.6°) | 23.9°<br>(20.9°-27.2°) |
|  | No-cursors | start level | 4.6°<br>(2.8°-6.9°) | 10.2°<br>(8.6°-11.9°) | 10.3°<br>(7.7°-12.8°) | 13.1°<br>(10.4°-16.4°) |

The fitted rate of change is relative to the asymptote. While in the first experiment (rotation size) these asymptotes vary with rotation size the absolute initial change (in degrees adaptation) in all four conditions are strikingly similar. Therefore, we report the absolute initial change as well.

### Exponential Decay of Washout

In our analysis of washout for the rotation size experiment, we employ an exponential decay function to investigate the decay of adaptation after the rotation has been removed. This function also used two parameters, the first denoted as R, representing the retention rate within the range of [1,1]. This indicates the proportion of the previous process value (P_t-1_) retained for the current trial (P_t_). Specifically, the process value at trial t (P_t_) is calculated by multiplying the value of the preceding trial (P_t-1_) by the retention rate (R). This multiplication operation reflects the diminishing effect of past information on the current state of the process. The second parameter of the function is the initial value of each process, denoted as P_t_ at the first trial (t=0), which falls within the range of [-1,2]*max(data). This range encapsulates the potential variability in initial values across different experimental conditions or datasets.

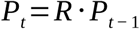

Both of the above equations describe the change of a quantity over time with parameters determining the rate of change or decay. However, the first equation models a process approaching an asymptote with a relative rate of change, while the second directly scales previous values with a retention rate, specific to washout decay. We again bootstrapped 5 k parameter values for each group, and the confidence intervals obtained from this are reported (see table 1). In order to compare parameters between groups, we subtract all the bootstrapped values from one group from all bootstrapped values of the other group, for 25 M difference scores. This is used to get a 95% confidence interval for the difference between groups. If this interval includes 0, we do not consider the difference to be meaningful in this data set.

### Step function

In the Continuous Aiming experiment we fit the emergence of explicit strategies in individual participants as a step function:

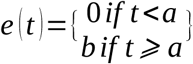

Such that *e*, the explicit strategy, is a function of the trial *t* . The strategy is 0 up to trial *a*, then steps up to the value *b*. In fitting this function, *a* and *b* are free parameters.

### Bayesian Statistics

We used Bayesian statistics to compare the extent of re-aiming during the rotated phase between different feedback types. Bayes Factors were used to determine whether there were significant differences or significant equivalences. Bayes Factors represent the ratio of the likelihood of the alternative hypothesis (the presence of a difference) to the likelihood of the null hypothesis (equivalence), given the data and a noninformative prior of √2/2. A Bayes Factor of 1 indicates an equal likelihood of both hypotheses. When the Bayes Factor falls within the range of 1/3 to 3, the evidence for both hypotheses is relatively equal, so this is considered anecdotal evidence with no strong preference for either hypothesis [22,23]. However, a Bayes Factor greater than 3 or less than 1/3 indicates moderate evidence in favour of the alternative hypothesis or the null hypothesis, respectively. Bayes Factors greater than 10 or less than 0.1 are considered strong evidence supporting one hypothesis over the other.

The approach was to first do a Bayesian “F-test” on the rate of change as well as the asymptotes across all groups within an experiment. If this indicated either little evidence one way or another or equivalence, no further test was done. If this indicated an effect of condition on either rate of change or asymptote, a series of Bayesian “t-tests” was done on only the parameters that showed an effect. In the rotation size experiment each pair of successively larger rotations was tested. In the feedback type and continuous aiming experiment, the control condition was compared with all other conditions. In the delayed feedback experiment, the terminal feedback condition without any delay was compared with all other conditions. In the continuous aiming experiment, there were only two conditions, such that no further tests were needed.

### Explicit adaptation

Using the 8 aiming trials in the second half of the rotated phase, we first test if the changed feedback affects the extent of explicit strategies as we’d expect. We also use this data to test the assumption of additivity of implicit and explicit adaptation, using two independent measures. The idea is that explicit and implicit adaptation add up to total adaptation. Even with variation in total adaptation, this should lead to a regression of implicit over explicit adaptation with a slope of −1 [5]. Hence, if the 95% confidence interval of the slope includes −1, we will consider this evidence for additivity.

## Results

To investigate potential changes in the extent and time-course of implicit adaptation in these groups, we compared rates of change (RofC) and asymptotes for both no-cursor trials and reach training trials. We also calculate means and 95% confidence intervals for the re-aiming responses. The resulting values, along with their bootstrapped 95% confidence intervals, are presented in Tables 1-4.

### Rotation Size

Previous results suggest that the extent of the implicit component of adaptation or aftereffects does not change with the size of the perturbation [6], but it is unknown whether the time-course is affected or not. We tested this for four rotations; 15°, 30°, 45°, and 60° (Fig 2 with N of 33, 54, 28 and 61, respectively) with a continuously visible cursor (for the outward reach) across different groups of participants. This experiment also contained a washout phase where the cursor-rotation was removed, so we could investigate how quickly people de-adapt both in no-cursor trials and reach-training trials (with cursor).

**Fig 2.**
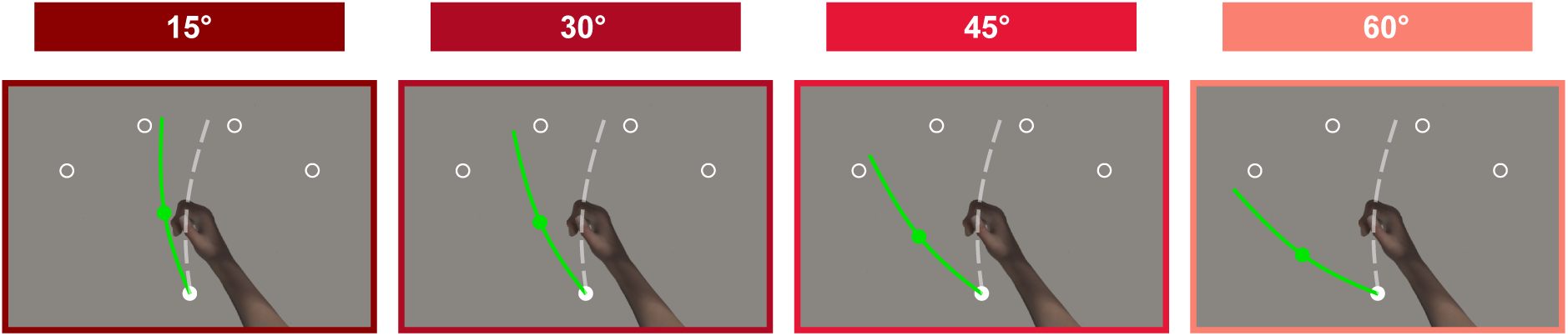
Participants reached with a cursor to one of the four forward targets as quickly and as straight as possible. Participants in the four conditions would train with one of four rotation sizes: 15°, 30°, 45° or 60° CCW rotated feedback.

Given that the magnitude of implicit aftereffects following adaptation typically remains relatively constant at around 15° regardless of the rotation size, we aim to investigate whether the time-course of these aftereffects is similarly unaffected by the magnitude of rotation. The results for cursor training and no-cursor reaches are plotted in Fig 3A and B respectively, and the fits are shown in Fig 3C-D. Statistical evaluation came from the 5 k parameter values for each group, which determined their confidence intervals (Table 1). We then calculated 25 million difference scores to compare groups and a 95% confidence interval for the group difference was obtained. If this interval includes 0, the difference is not considered significant in this dataset. First, we look at reach-training trials during the longer learning phase for all successive rotation sizes. We find that rates of change are not different [95% CI for 15° to 30° (−0.075, 0.085); 30° to 45° (−0.095, 0.097); 45° to 60° (−0.041, 0.144)], but asymptotes increase with rotation size [95% CI for 15° to 30° (−10.474, −6.815); 30° to 45° (−17.167, −9.392); 45° to 60° (−14.433, −5.499)], as expected (Table 1). For no-cursors, the 30° group at 18.5% (14.6%-23.2%) has a larger RofC than the 45° group at 11.0% (7.9%-16.5%), meaning that people training with the 30° rotation change their no-cursor reach directions faster than the 45° group, as seen in Table 1 (95% CI 0.009, 0.132). Otherwise, there are no differences in learning rates in the comparisons here [95% CI for 15° to 30° (−0.035, 0.146); 45° to 60° (−0.016, 0.079)].

**Fig 3.**
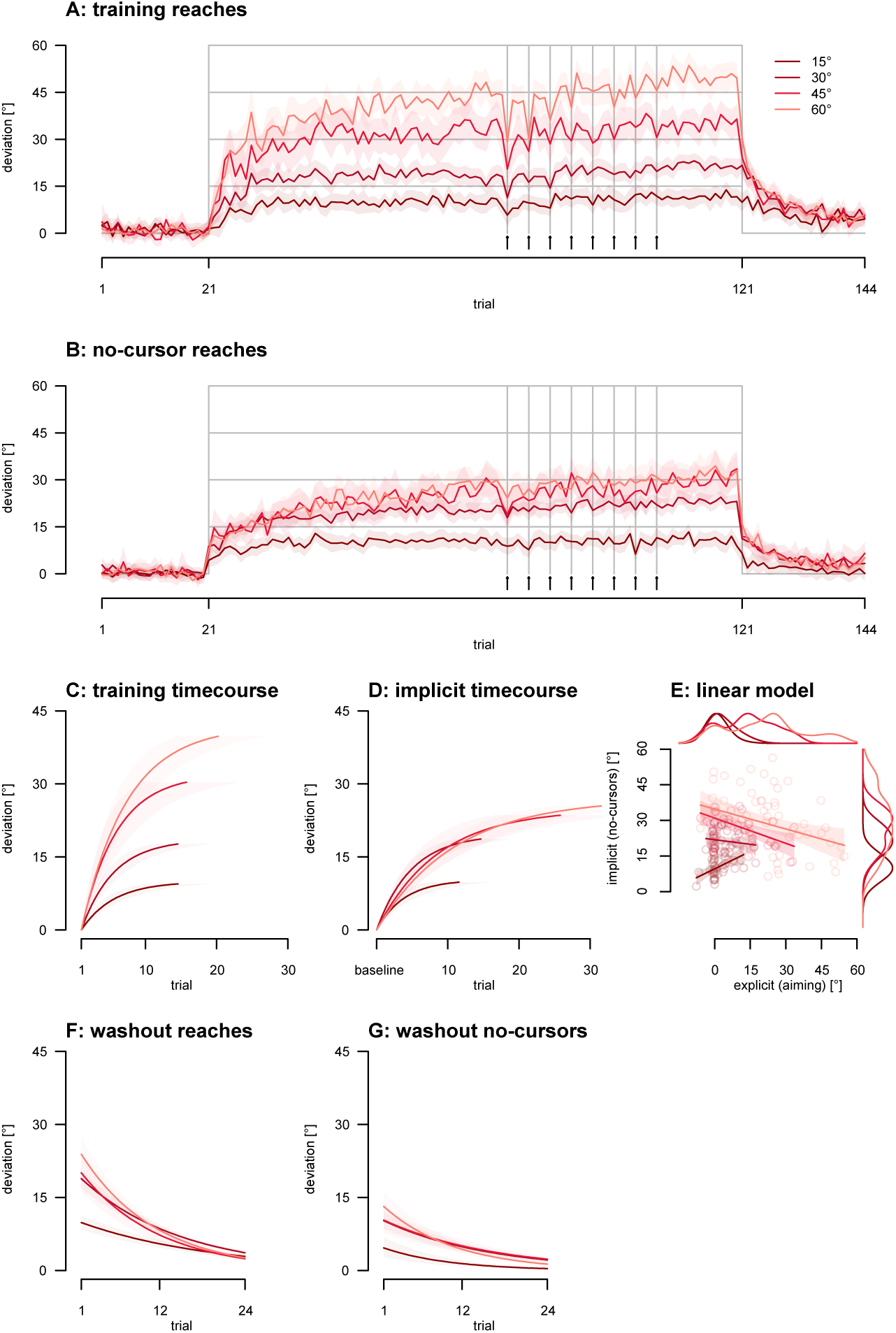
Rotation Size. Shaded regions indicate 95% confidence intervals. **A.** Reach adaptation across trials, with eight aiming trials in the second half of the aligned phase (indicated by arrows and vertical lines). **B.** Implicit reach aftereffects across trials. **C.** Fitted exponential curves for reach adaptation in the rotated phase. **D.** Fitted exponential curves for implicit reach aftereffects in the rotated phase. **E.** Individual data scatter plot with regression lines depicting the relationship between implicit and explicit learning processes. Each dot represents a participant. **F.** Fitted exponential decay for reach training trials in the washout phase. **G.** Fitted exponential decay for reach aftereffects in the washout phase.

**Fig 4.**
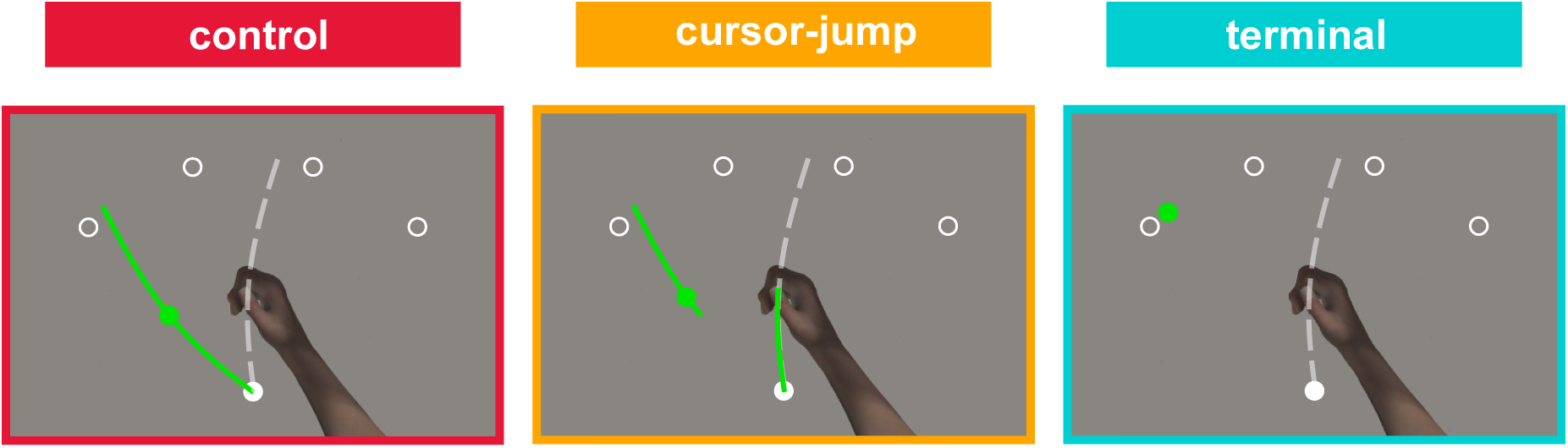
Participants reached with a cursor to one of the four or eight forward targets as quickly and as straight as possible. Participants in these three groups experience different trial types through three kinds of rotated cursor feedback: continuous, terminal (cursor only shown at end of reach trial), and cursor jump (cursor jumps 45° CCW mid-reach on every trial). Each participant would perform a trial with one of the above feedback types interleaved with no cursor trial.

Surprisingly, we found that the implicit aftereffects were not simply capped at 15° but varied slightly with rotation size, as shown in Fig 3B. Specifically, implicit asymptotes did vary across rotations of 15°, 30°, and 45° [95% CI for 15° to 30° (−11.101, −7.476); 30° to 45° (−9.114, −1.435)]. Only when the rotation increased from 45° to 60° [95% CI for 45° to 60° (−6.569, 2.406)] did we fail to detect a difference in our large sample sizes.

Now we look at how these groups compare during washout and the fits for these washout trials are shown in Fig 3 F&G. We compare the 15° group reach training parameters with those from all other groups and find that rates of change did not differ between the 15° and 30° groups [95% CI for 15° to 30° (−0.007, 0.039)], but the washout in the 15° group is slower than in the 45° and 60° group [15° to 45° (0.010, 0.062); 15° to 60° (0.015, 0.064)]. Additionally, the 15° group has a smaller starting level (Table 1, and left side of Fig 3B) then all other groups [95% CI for 15° to 30° (−11.312, −6.667); 15° to 45° (−13.994, −6.193); 15° to 60° (−17.628, −10.687)] whose start levels seem to overlap each other. The same pattern is observed when examining no-cursor rates of change [95% CI for 15° to 30° (−0.190, 0.026); 15° to 45° (−0.198, 0.038); 15° to 60° (−0.162, 0.062)] and starting levels [95% CI for 15° to 30° (−8.103, −2.806); 15° to 45° (−8.815, −2.292); 15° to 60° (−12.179, −4.911)].

Here we will investigate explicit strategy as assessed by the eight aiming trials. We expect little explicit strategy in the 15° and 30° conditions, and will see how this develops in larger rotations. We find an effect of rotation size on the magnitude of re-aiming responses (BF_10_ > 1000, as illustrated by the density plots along the x-axis in Fig 3E and the first 4 bars in Fig 11E). We then compared the re-aiming responses to 0 for each condition, and then completed follow-ups between successive rotations. There is no evidence for or against re-aiming in the 15° condition (0.9° on average, BF_10_ = 0.489), however in the other three conditions, it is much more clear that almost all participants engage in some amount of re-aiming (BF_10_ > 1000, also seen in Fig 11E). Now comparing aiming between successive rotations, we find that re-aiming is moderately different between the 15° and 30° groups (BF_10_ = 5.115). We do see the expected difference between the 30° and 45° group (BF_10_ > 1000), but there is no evidence when comparing the 45° and 60° group (BF_10_ = 1.959). For the most part, we observe comparable explicit changes in the two largest rotations.

Finally, we will explore how explicit adaptation fares as a predictor of implicit adaptation (Fig 3E). The linear relationship is significant in the 15° group, but the confidence interval for the slope does not include −1 (which would indicate additivity) [r = 0.461, p = 0.007, slope: 0.488 CI (0.144, 0.832)]. The confidence interval of the slope also does not include 0 (no relationship) and has positive values, so there is no evidence for additivity of implicit and explicit adaptation. The 30° group had no significance [r = −0.121, p = 0.381, slope: −0.129 CI (−0.421, 0.164)]. For the 45° and 60° groups, the linear relationship is significant, but the confidence interval for the slope does not include −1 [45°: r = −0.539, p = 0.004, slope: −0.356 CI (−0.568, −0.127); 60°: r = −0.356, p = 0.005, slope: −0.279 CI (−0.469, −0.088)]. Taken together, we find a weak and non-additive relationship between implicit and explicit adaptation.

### Feedback Type

We tested three different types of feedback in the Feedback-Type experiment: Continuous, Terminal and Cursor Jump (Fig 4). There were 51 participants in the *Continuous-feedback or Control Group,* the cursor was continuously visible during the outward cursor movement. This included the 24 participants from the previous experiment who did the 45° condition. The 35 participants who were in the *Terminal Group* were only shown the rotated cursor at the end of the outward movement. Specifically, the cursor was not displayed until the hand had moved 8.8 cm radially from home position. There was also only 1 static cursor position shown for 750 ms, regardless of any subsequent movements by the participant. That is, visual feedback consisted of knowledge of outcome only. In line with this, the return to the home position was guided by the same green circle feedback as used in the no-cursor reaches. The *Cursor Jump Group* consisted of 32 participants and the cursor for this group was aligned with the hand for the first half of the distance to the target. When this 50% distance was reached, the cursor rotation of 45° (CCW) was applied for the rest of the trial. This type of feedback was similar to an earlier task from our lab [15] which increased explicit strategy.

In this experiment, we wanted to understand how rapidly the implicit components of adaptation emerge in response to the type of visual information during classical visuomotor adaptation. The results for cursor training and no-cursor reaches are plotted in Fig 5 A and B respectively, and the fits are shown in Fig 5C and D. From our bootstrapped parameters, we computed confidence intervals for each group (Table 2). We then compared the groups by calculating difference scores and deriving a 95% confidence interval for their difference. If this interval includes 0, the difference is not deemed significant in this dataset. First, looking at reach training in Fig 5, it seems that the three groups have similar rates of change, and this is confirmed by the 95% confidence intervals for the difference between groups [control and terminal (−0.276, 0.025); control and cursor jump (−0.006, 0.161)]. Similarity is also observed for reach training asymptotes [control and terminal (−4.393, 3.172); control and cursor jump (−6.804, 1.478)]. We now compare the test groups with the control group on parameters describing the time-course of implicit reach aftereffects. We find no effect on rates of change [control and terminal (−0.057, 0.080); control and cursor jump (−0.066, 0.053)], but the asymptotic level of implicit reach aftereffects is larger for the control group than for either test group [control and terminal (5.160, 13.296); control and cursor jump (2.482, 10.623)], with the control group at 24° (21.7°-26.5°) and terminal and cursor jump having asymptotic values of 14.6° (11.7°-18.3°) and 17.4° (14.3°-21.0°), respectively.

**Fig 5.**
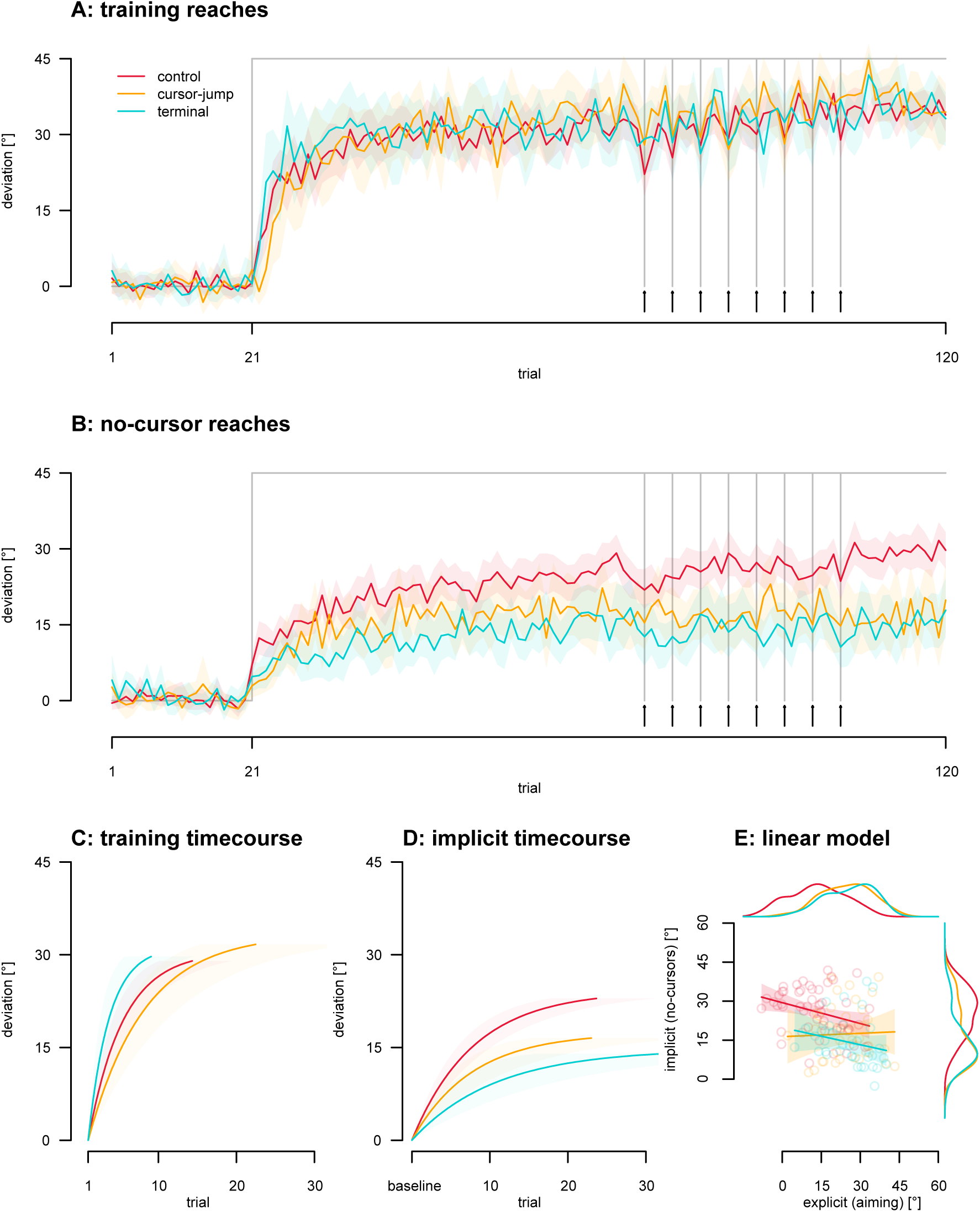
Feedback Type. Shaded regions indicate 95% confidence intervals. **A.** Reach adaptation across trials, with eight aiming trials near the end of the rotated phase (indicated by vertical lines and arrows). **B.** Implicit reach aftereffects across trials **C.** Fitted exponential functions for reach adaptation in the rotated phase. **D.** Fitted exponential functions for implicit reach aftereffects in the rotated phase. **E.** Scatter plot with regression lines depicting the relationship between implicit and explicit learning processes. Each dot represents a participant.

**Table 2.** Descriptives of adaptation for each group in the feedback type experiment. Rates of change (RofC) and asymptotes are shown for training and no-cursor reaches in the rotated phase. Aiming extent is shown as the average across the eight aiming trials in the second half of the rotated phase. The entries for RofC and asymptote both first list (what could be considered) the group average using each participant once, then in parentheses the 95% confidence interval of the mean, based on 5k bootstraps across participants. For RofC we also list the absolute value after one rotated trial as predicted by the function fitted to all data.

|  |  | control (N=58) | cursor-jump (N=33) | terminal (N=35) |
| --- | --- | --- | --- | --- |
| Reach training | RofC | 20.1%<br>(15.0%-28.0%)<br>6.1° | 12.9%<br>8.4%-18.6%)<br>4.3° | 30.9%<br>(19.9%-46.9%)<br>9.6° |
|  | asymptote | 30.5°<br>(27.9°-33.2°) | 33.1°<br>(30.0°-36.3°) | 31.2°<br>(28.4°-33.8°) |
| No-cursors | RofC | 11.9%<br>(9.2%-15.5%)<br>2.9° | 12.2%<br>(7.9%-17.8%)<br>2.1° | 9.0%<br>(5.4%-17.3%)<br>1.3° |
|  | asymptote | 24.0°<br>(21.7°-26.5°) | 17.4°<br>(14.3°-21.0°) | 14.6°<br>(11.7°-18.3°) |
| Re-aiming | extent | 13.3°<br>(10.6°-16.0°) | 24.7°<br>(21.4°-27.8°) | 25.8°<br>(22.8°-28.9°) |

Testing re-aiming, we find an effect of feedback type on explicit strategies (BF_10_ > 1000, as seen in Fig 9E). We can see in Table 2 and Fig 5 that terminal and cursor jump have 170% the amount of explicit strategy as compared to the control group (∼14° vs. ∼25° of strategy), and comparing the control group to the others we find differences from terminal and cursor jump (both BF_10_ > 1000).

Here we test explicit adaptation as a predictor of implicit adaptation. Across participants in the control group there is a significant relationship, but the confidence interval for the slope does not include −1 [r=-0.321, p=0.016, slope: −0.268 CI (−0.484, −0.053)], as illustrated in Fig 5E. This matches the pattern found before in the rotation size experiment (Fig 3E). However, none of the other relations are significant, and the confidence intervals of the slopes do not include −1 (which would indicate additivity) but they do include 0 (i.e. no relationship) [terminal: r=-0.213, p=0.219, slope: −0.220 CI (−0.577, 0.137); cursor jump: r=0.038, p=0.834, slope: 0.044 CI (−0.378, 0.465)]. Thus, there is only a weak relationship between implicit and explicit components in the control group, but none for the terminal and cursor jump conditions.

### Feedback Delay

After testing the kind of feedback, we now wanted to check how manipulating the timing of feedback affected implicit learning (Fig 6). Studies have claimed that inserting a delay prior to terminal feedback can reduce the extent of implicit learning [9,16]. However, the impact of this delay on the rate of implicit learning remains unknown. The following experiment seeks to shed light on this.

**Fig 6.**
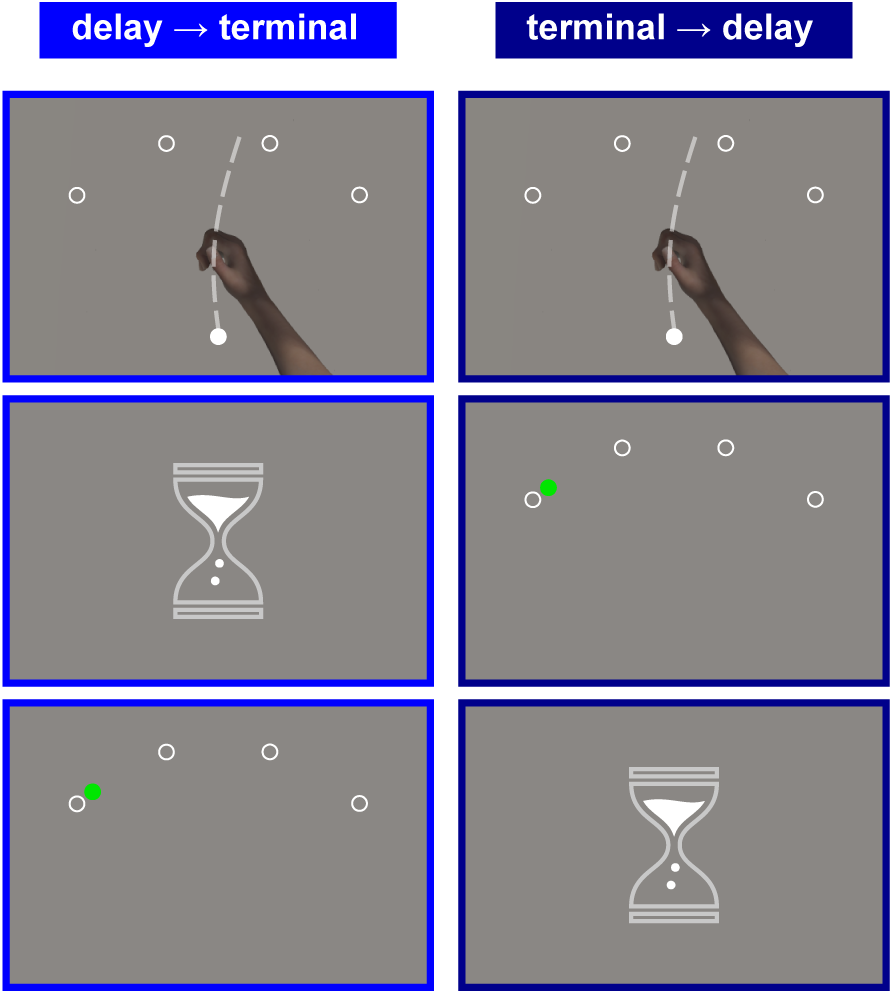
Participants in the two new groups in this experiment experienced a delay as follows. The delay → terminal group would make the reach, wait during the 1.2 s delay, and then receive feedback about the end-point position for 0.6 s. The terminal → delay would receive feedback right away for 0.6 s, and then wait for 1.2 s. The delay and feedback intervals combined took 1.8 s in each case, and the participant was to hold the stylus during that time, and only return to the home position after the 1.8 s hold. Each participant would perform a trial with one of the above types of feedback interlaced with a no cursor after every trial.

### delay → terminal group

26 participants adapted to a 45° CCW visuomotor rotation with terminal cursor feedback. This group was like the Terminal Group in the feedback type experiment, except that we included a 1.2-second delay before the cursor was displayed for 600 ms following the 8.8 cm outward hand movement. After participants had moved their hand 8.8 cm away from the start position and held their final position for 1.2 s, they received end-point position feedback for 600 ms. Participants were instructed to hold their end position for the full 1.8 seconds that took up the delay and the feedback period. After participants had held their end position for 1.8 seconds, the green circle guiding return movements would be shown, signalling that participants could move back to the home position. However, if participants moved more than 0.4 cm away from the end position of their reach during the hold period, they would need to move to any point, at least 8.8 cm away from home again, and restart the hold period of 1.8 seconds. This was to motivate participants to indeed hold the stylus at the end position while feedback was shown. The terminal feedback was shown at the same time and for the same duration after the outward reach was finished, irrespective of whether or not the hold was maintained.

### terminal → delay group

The 39 participants in this group served as a control for the above delay → terminal group in case the extra delay time would affect the overall time-course. For this group, much like the original Terminal group, they received a single position of cursor feedback for 600 ms immediately after the pen moved to the edge of the outer stencil where the targets were displayed. But we inserted a 1.2-second delay afterwards so that the trial length was identical to that in the delay → terminal group, which controls for the increased inter-reach interval. Like the delay → terminal group, this group also had to maintain a 1.8-second hold at the end of the reach, and had to restart the hold if it was broken before 1.8 seconds had elapsed. These two groups had the same durations of trials and the whole experiment, with the only difference in the timing of the feedback. The terminal → delay group immediately got the feedback, but the delay → terminal group had to wait 1.2 s after reach completion before seeing the terminal feedback. According to previous studies [9,14,16], this should lead to more explicit adaptation, and hence perhaps would also lead to less, or slower, implicit adaptation. We compare these two experimental groups with the previous terminal group that had no delays, as well as with the overall control group, as illustrated in Fig 7.

**Fig 7.**
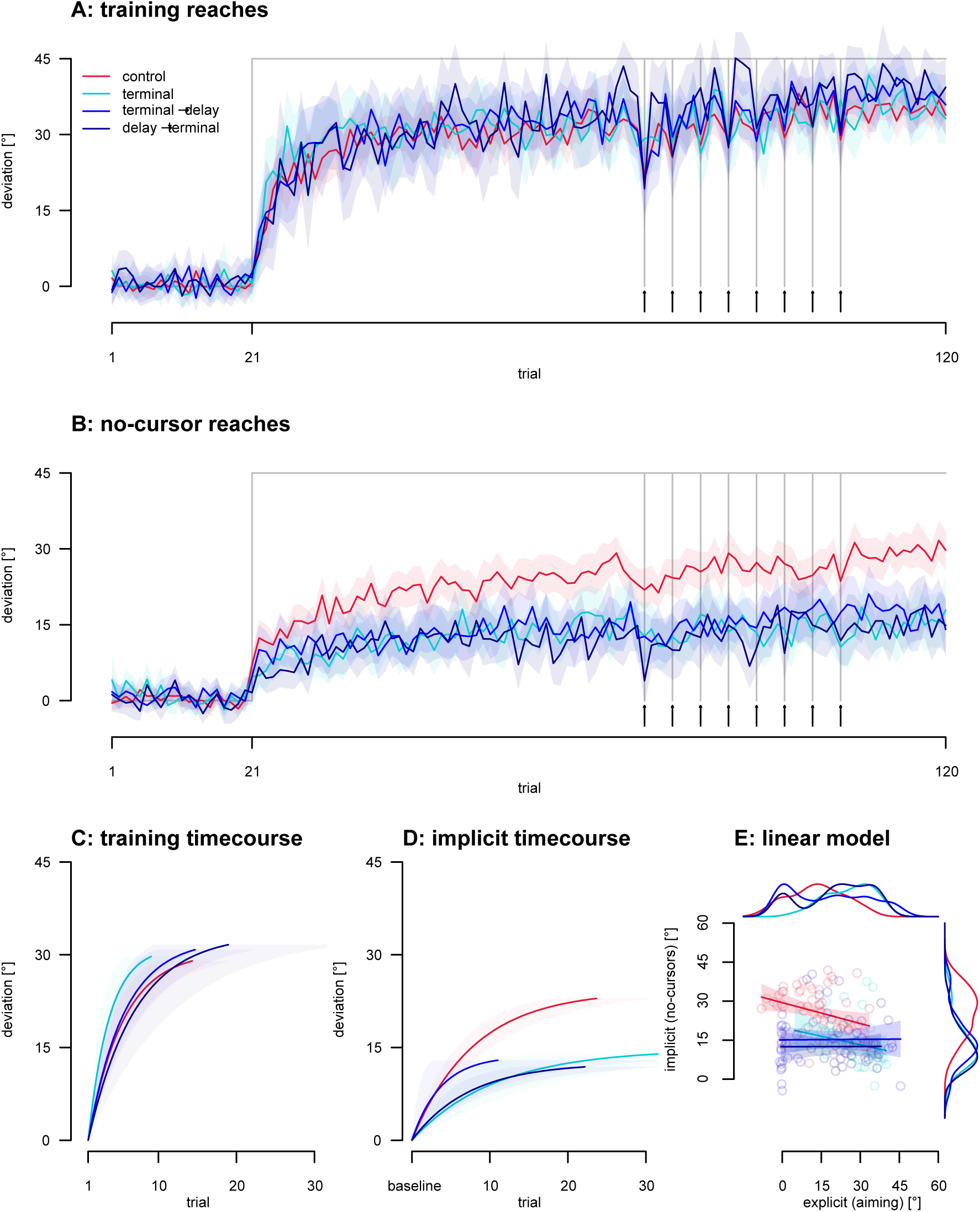
Feedback Delay. Shaded regions indicate 95% confidence intervals. **A.** Reach adaptation across trials, with eight aiming trials in the rotated phase indicated by arrows and vertical lines. **B.** Implicit reach aftereffects across trials **C.** Fitted exponential functions for reach adaptation in the rotated phase. **D.** Fitted exponential functions for implicit reach aftereffects in the rotated phase. **E.** Scatter plot with regression lines depicting the relationship between implicit and explicit learning processes. Each dot represents a participant.

Seeing that the feedback type experiment found an effect of terminal feedback on the asymptote of implicit reach aftereffects, this experiment asks if there are any additional effects of delays when combined with terminal feedback. The results for cursor training and no-cursor reaches for these conditions are plotted in Fig 7A and B respectively, and the fits are shown in Fig 7C-D. At first glance (Fig 7; Table 3) the two terminal feedback groups with delays look very similar to the terminal feedback group without delays. We compare the two new groups to the previous terminal group, using 95% confidence intervals from difference scores. We find no effect on overall adaptation for both RofC [95% CI for terminal and delay → terminal (−0.017, 0.329); terminal and terminal → delay group (−0.033, 0.287)], and asymptotes [95% CI for terminal and delay → terminal (−6.402, 2.378); terminal and terminal → delay group (−4.967, 2.433)] (Fig 7, Table 3). As illustrated in Fig 7B, we find a similar absence of a difference for the implicit no-cursors across the different terminal conditions (blue curves), for both RofC [95% CI for terminal and delay → terminal (−0.113, 0.053); terminal and terminal → delay group (−0.564, 0.028)] and asymptote [95% CI for terminal and delay → terminal (−2.202, 6.686); terminal and terminal → delay group (−3.312, 5.558)], as shown in Fig 7 and Table 3. In summary, adding a delay before or after the cursor feedback did not reduce rates of change or asymptotes in either reach training or implicit reach aftereffects.

**Table 3.** Descriptives of adaptation for each group in the feedback delay experiment. Rates of change (RofC) and asymptotes are shown for training and no-cursor reaches in the rotated phase. Aiming extent is shown as the average across the eight aiming trials in the second half of the rotated phase. The entries for RofC and asymptote both first list (what could be considered) the group average using each participant once, then in parentheses the 95% confidence interval of the mean, based on 5k bootstraps across participants. For RofC we also list the absolute value after one rotated trial as predicted by the function fitted to all data.

|  |  | delay → terminal<br>(N=28) | terminal → delay<br>(N=39) | terminal (N=35) | control (N=58) |
| --- | --- | --- | --- | --- | --- |
| Reach training | RofC | 15.4%<br>(8.6%-29.8%)<br>5.1° | 19.5%<br>(13.1%-29.6%)<br>6.3° | 30.9%<br>(19.9%-46.9%)<br>9.6° | 20.1%<br>(15.0%-28.0%)<br>6.1° |
|  | asymptote | 33.1°<br>(29.7°-36.6°) | 32.3°<br>(29.9°-35.0°) | 31.2°<br>(28.4°-33.8°) | 30.5°<br>(27.9°-33.2°) |
| No-cursors | RofC | 12.6%<br>(8.8%-19.5%)<br>1.6° | 23.9%<br>(8.7%-65.5%) 3.2° | 9.0%<br>(5.4%-17.3%)<br>1.3° | 11.9%<br>(9.2%-15.5%)<br>2.9° |
|  | asymptote | 12.5°<br>(9.7°-15.7°) | 13.5°<br>(10.9°-16.8°) | 14.6°<br>(11.7°-18.3°) | 24.0°<br>(21.7°-26.5°) |
| Re-aiming | extent | 20.9°<br>(16.0°-25.5°) | 17.9°<br>(13.2°-22.4°) | 25.8°<br>(22.7°-28.8°) | 13.3°<br>(10.8°-16.1°) |

**Table 4.**
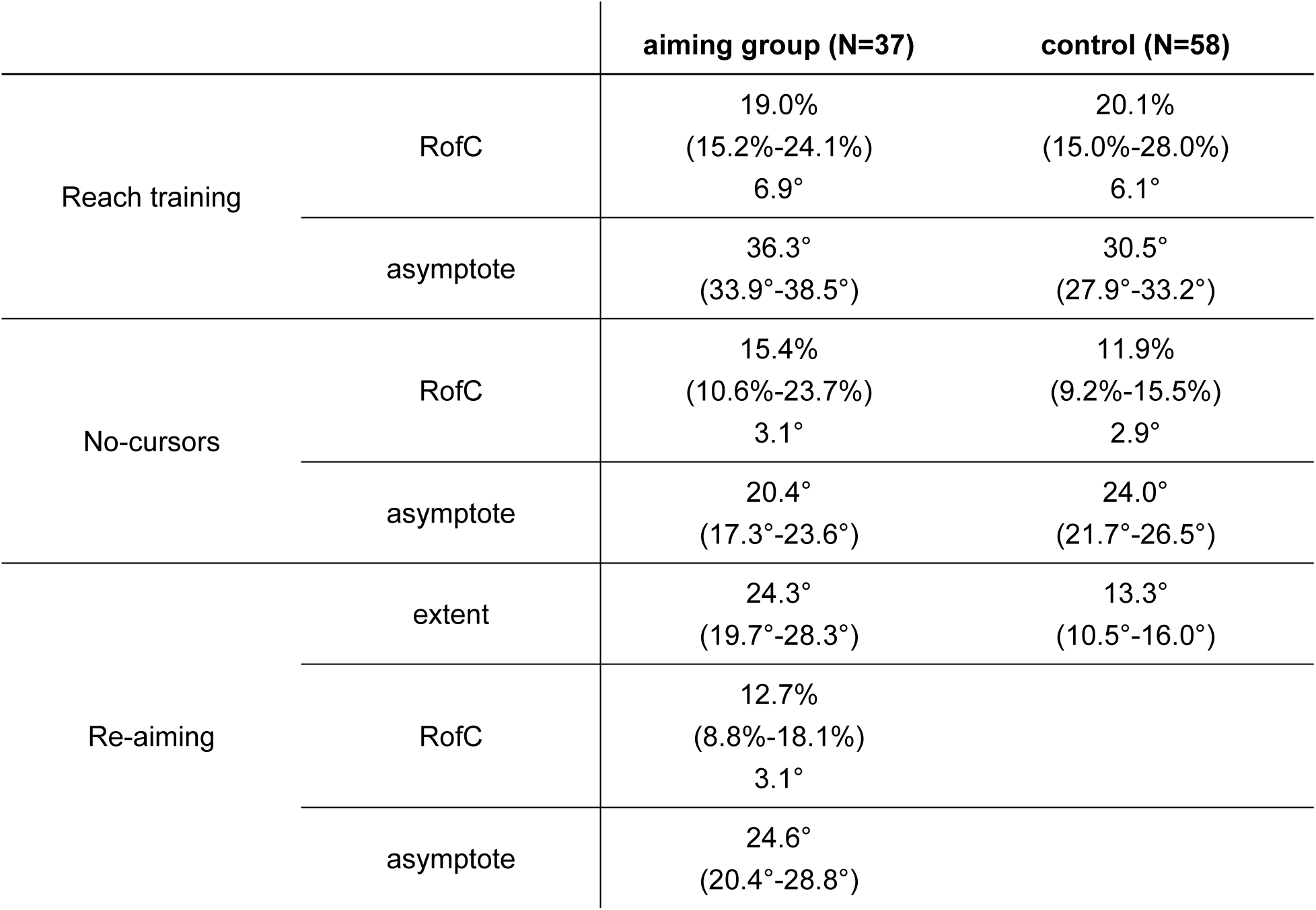
Descriptives of adaptation for each group in the continuous aiming experiment. Rates of change (RofC) and asymptotes are shown for training and no-cursor reaches in the rotated phase in both groups, and for the re-aiming responses in the continuous aiming group. Aiming extent is also shown as the average across the eight aiming trials in the second half of the rotated phase (using the same 8 trials for both groups). The entries for RofC and asymptote both first list (what could be considered) the group average using each participant once, then in parentheses the 95% confidence interval of the mean, based on 5k bootstraps across participants. For RofC we also list the absolute value after one rotated trial as predicted by the function fitted to all data.

Then, we investigated if there was an effect of delays on aiming, and interestingly we do see one (BF_10_ = 546), as seen in Fig 11E. Follow up tests show there is little evidence for either a difference or equivalence between the terminal group and the delay → terminal group (BF_10_ = 0.915), but the re-aiming responses are smaller in the terminal → delay group as compared to the terminal feedback group (BF_10_ = 4.689). That is: adding a delay after the terminal feedback seems to make adaptation *less* explicit (Fig 9).

Finally, we test explicit adaptation as a predictor of implicit adaptation. The two groups with delays again show no linear relationship between implicit and explicit adaptation [delay → terminal: r=0.004, p=0.983, slope: 0.003 CI (−0.234, 0.239); terminal → delay: r=-0.010, p=0.952, slope: 0.007 CI (−0.218, 0.232)].

### Continuous Aiming

Previous work from our lab showed that aiming trials throughout a visuomotor adaptation paradigm can lead to more explicit adaptation [5]. This is why we avoided using aiming trials until after adaptation was close to saturation. However, to test if this assumption is true and to see if explicit adaptation is indeed faster than implicit adaptation, we also included a continuous aiming condition. Instead of participants conducting 8 aiming trials late into the rotated phase like in all previous experiments to measure explicit strategy, this condition introduced consistent aiming trials throughout the rotated phase (Fig 1E). Participants performed an aiming trial, followed by a reach-training trial and a no-cursor trial in a repeated pattern. Thus, we had a single experimental group in the continuous aiming version of the experiment that adapted to a 45° rotation to examine the time-course of both explicit strategy use and development, and implicit adaptation. The results of these two conditions, including the aiming (dashed lines), across trials are shown in Fig 8 A&B, with fits plotted in Fig 8C&D.

**Fig 8.**
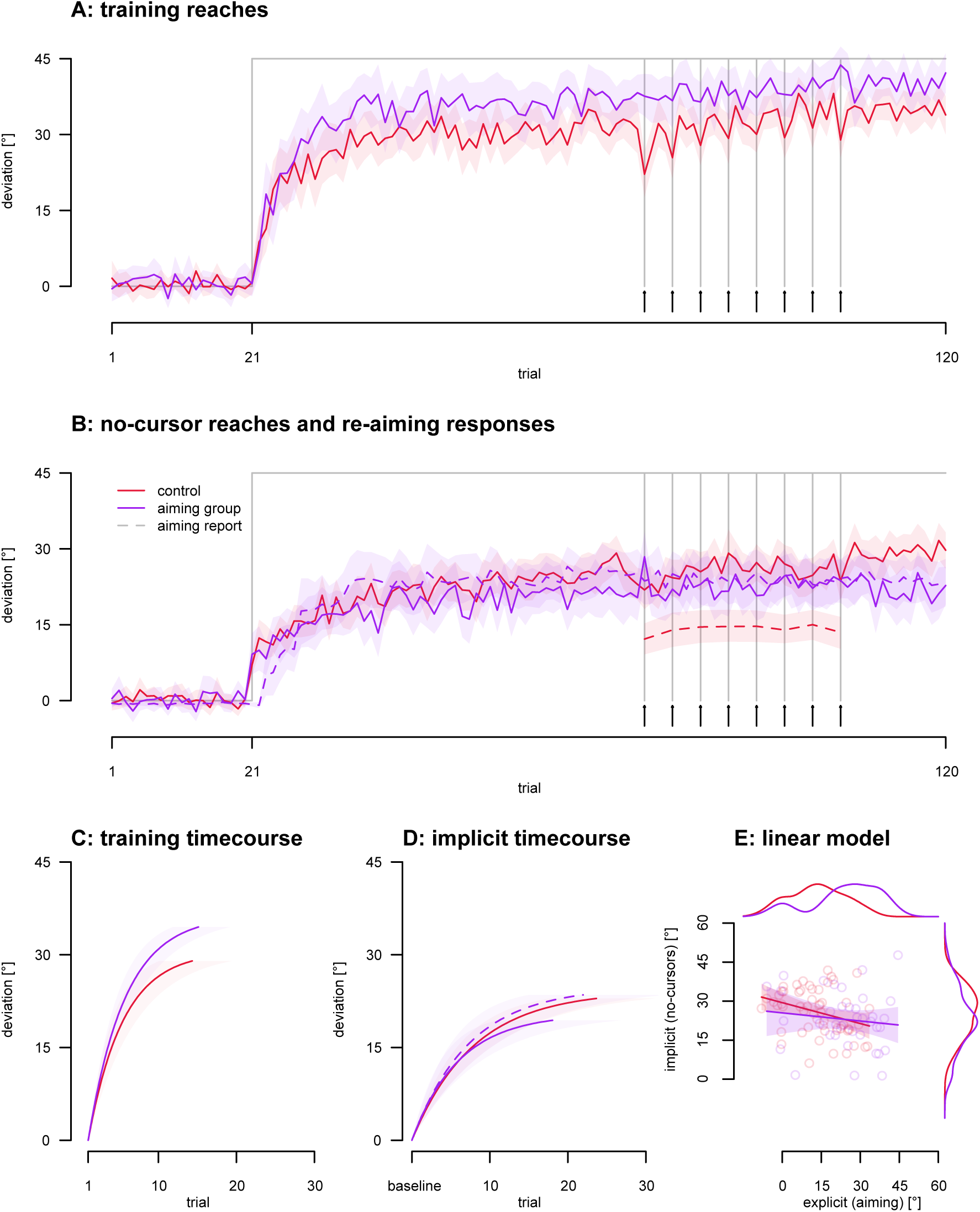
Continuous Aiming. Shaded regions denote 95% confidence intervals. **A.** Reach adaptation across trials, with eight aiming trials indicated by vertical lines and arrows. **B.** Implicit reach aftereffects across trials (solid lines) and explicit re-aiming for the continuous group (purple dashed lines) and the control condition (red dashed line). **C.** Rate of Change for rotated phase of Reach adaptation **D.** Fitted exponential curve for implicit reach aftereffects in the rotated phase, and for the explicit re-aiming in the continuous group (purple dashed curve). **E.** Scatter plot with regression lines depicting the relationship between implicit and explicit learning processes. Each dot represents a participant.

Bootstrapping our parameters, we derived confidence intervals for each group (Table 4). Then, by calculating difference scores, we obtained a 95% confidence interval to compare the groups. If this interval includes 0, the difference is not considered significant in this dataset. At first glance (Fig 8A-E; Table 4), there might be larger adaptation in the continuous aiming group, with no clear effects on the amount of implicit adaptation or the rates of change of both reach training and no-cursor reaches. Indeed, the new continuous aiming group has an increased extent of overall adaptation [95% CI (−9.229, −2.175)], with no clear effect on the extent of implicit adaptation [95% CI (−0.269, 7.582)]. There are no differences in the rate of overall adaptation [95% CI (−0.061, 0.097)] and the rate of implicit adaptation [95% CI (−0.269 −7.582)].

Additionally, in this group, we were interested in comparing the rate of change of aiming responses (purple dashed lines in Fig 8B&D) with the rate of change of implicit adaptation. This tests the assumption that implicit adaptation is slower than explicit adaptation. We find no differences in the RofC of explicit cognitive strategy (dashed purple lines in Fig 8B&D) compared to implicit no cursor RofC (solid lines in Fig 8B&D) in the control group [95% CI (−0.069, 0.044)] or in the continuous aiming group [95% CI (−0.046, 0.117)]. Specifically, as listed in Table 4, for the continuous aiming group, the RofC for explicit re-aiming was 12.7% (8.8%-18.1%) and 15.4% (10.6%-23.7%) for the implicit reach aftereffects; this led to a fitted change in deviation after the first training trial of 3.1° for both implicit and explicit measures. Thus, with the current data set and approach, we can not detect a difference in how quickly implicit or explicit adaptation emerges, so it is possible they might be equally fast in this group. That is, implicit and explicit contributions to adaptation emerged simultaneously and at the same rate in this data set.

For our next analysis, we wanted to see if there is a difference in the reported aiming direction between the continuous aiming group and the control group. Analyzing the 8 aiming trials during the latter portion of the rotated phase, we compare the control group of participants performing aiming trials only 8 times (red dashed line in Fig 8B) with those that do aiming trials throughout the whole experiment (purple dashed line in Fig 8B, see Fig 11E too). Our findings indicate a difference in explicit strategy between these participant groups (BF_10_ = 767), which is aligned with the 95% confidence intervals in Table 4, which do not overlap. Notice however that strategies are not really normally distributed, but tend to cluster at specific magnitudes (figure 3E, 5E, 7E and 8E). Moreover, while adding continuous aiming may evoke strategies in some participants who would not have discovered one themselves, a majority of participants in the control group did have a strategy already.

Lastly, we test explicit adaptation as a predictor of implicit adaptation for the aiming group. There seems to be a significant, but non-additive relationship between measures of implicit and measures of explicit adaptation (r=-0.321, p=0.016, slope: −0.268 CI (−0.484, −0.053)] as illustrated in Fig 8E. In other words, consistent with the results above, the implicit reach aftereffect did not consistently vary with the magnitude of explicit strategies, as should be the case if the implicit component were merely the residual difference between overall adaptation and the strategy used.

### Subtractive estimate of implicit adaptation

Some recent work uses re-aiming responses subtracted from total adaptation as an estimate of implicit adaptation [18,19]. This relies on additivity being true, and our data sets suggest that this is unlikely. Nevertheless, we investigate the time course of this estimate to compare it to the other processes we measured, and to allow a comparison with other studies. Our expectation was that subtractive estimates of implicit adaptation would be slow, given their characterization as the slow process [17].

First, we look at the data in Fig 9A and see that the subtractive measure increases sharply during the second rotated trial. This jump likely results from some adaptation in the training reaches, while conversely, the aiming responses adjust more slowly. Also, the no-cursor reaches increase when the rotation is introduced, due to participants having already encountered a rotated trial in the training phase. As mentioned before, this prior experience is accounted for in our fits by adding a zero at the start of the time series.

Comparing the rate of change of the subtractive estimate of implicit adaptation (Fig 9B), we find that it surpasses all other types of data time courses in speed [95% CI for subtraction to reaches (0.009, 0.683); subtraction to aiming (0.072, 0.746); subtraction to no-cursors (0.038, 0.719); see Table 5]. Specifically, the fitted time courses show that the subtractive measure reaches 95% of its asymptote by the second trial (although with a wide confidence interval), marking it as the fastest among the group. Additionally, the subtraction approach yields a smaller asymptote compared to direct estimates from implicit no-cursors [95% CI for subtraction to no-cursors (−10.761, −1.204), also shown in Table 5]. Essentially, implicit adaptation is faster than anything else if estimated this way. This is not what we expected to see. And, even though we have shown before that implicit adaptation can be fast in some cases, this result seems highly unlikely to us. It should for now perhaps be seen as a peculiarity.

**Table 5.** Descriptives of each data type compared to subtraction in our experiments. Rates of change (RofC) and asymptotes are shown for reaches, no-cursor reaches, re-aiming trials, and subtraction in the rotated phase. The entries for RofC and asymptote both list (what could be considered) the group average using each participant once, then in parentheses the 95% confidence interval of the mean, based on 5k bootstraps across participants. For RofC we also list the absolute value after one rotated trial as predicted by the function fitted to all data.

|  | reaches | re-aiming | no-cursors | subtraction |
| --- | --- | --- | --- | --- |
| RofC | 19.0%<br>(15.2%-24.1%)<br>6.9° | 15.4%<br>(10.6%-23.7%)<br>3.1° | 12.7%<br>(8.8%-18.1%)<br>3.1° | 64.3%<br>(20.7%-87.6%)<br>7.8° |
| asymptote | 36.3°<br>(33.9°-38.5°) | 20.4°<br>(17.3°-23.6°) | 24.6°<br>(20.4°-28.8°) | 12.1°<br>(10.9°-18.1°) |

Next, we did regressions on the direct, no-cursor measures over the subtractive estimates of implicit adaptation when processes are close to saturation (Fig 9D). They show that these two variables only share a small amount of variance, and this holds across the continuous aiming group and all other groups undergoing a 45° rotation [continuous aiming: r=0.080, p=0.639, slope: 0.060 CI (−0.198, 0.319); all 45° groups: r=-0.145, p=0.045, slope: 0.123 CI (0.003, 0.243)]. This might not be surprising given that implicit measures over explicit measures, show roughly the same relationship [continuous aiming: r=−0.124, p=0.466, slope: −0.105 CI (−0.392, 0.183); all 45° groups: r=−0.226, p=0.002, slope: −0.193 CI (−0.313, −0.073), shown in Fig 9C]. We should and do see an equivalent relationship between predicted implicit adaptation using the subtractive method and directly measured implicit adaptation, except with the direction flipped. It appears there is no clear relationship between measures of explicit and implicit adaptation in this data set, whether these measures are subtractive or direct.

### Nature of explicit time course

The time course of subtractive estimates relies on 2 other time courses as well as an assumption of additivity. This further relies on the nature of the timecourses of implicit and explicit adaptation being comparable. At first glance, the average timecourses do look similar, (Fig 9A) but is the average a good representation of the process? While levels of implicit adaptation seem to fall in a more uni-modal distribution, aiming reports show two clusters of responses: a small one at 0° (for no strategy) and another, larger one at roughly 20-40° (see Fig 8E) which are more clearly separated in the continuous aiming group.

To understand where these clusters of strategies come from, we look at the individual timecourses for overall, implicit and explicit adaptation (Fig 10). While noisy, it seems that for individual participants, both overall and implicit adaptation gradually increases (Fig 10A&C). On the other hand, all participants who shift their strategy from 0 to the second cluster do so in a fairly discrete step within the first 30 trials, most of them within the first 10 trials (Fig 10E). Since they do this at various times, the average time course of aiming responses still looks like an exponential learning curve. To test if explicit adaptation is better characterized as sudden moments of insight, perhaps akin to aha!-learning, we fit exponential functions and step functions to the individual time course of overall, implicit and explicit adaptation for each participant. We then compare the RMSEs between the fit and the data for each participant, using paired, Bayesian t-tests (Fig 10B,D,F). While the exponential functions seem to fit overall adaptation slightly better, there is no evidence for this (BF_10_=2.4). There is evidence that implicit adaptation is better fit by exponential functions (BF_10_=10.7) and explicit adaptation is better fit by step functions (BF_10_=7.3). This shows a difference in how implicit and explicit adaptation emerge, gradually or suddenly, indicating that the two processes may be of a very different nature.

**Fig 9.**
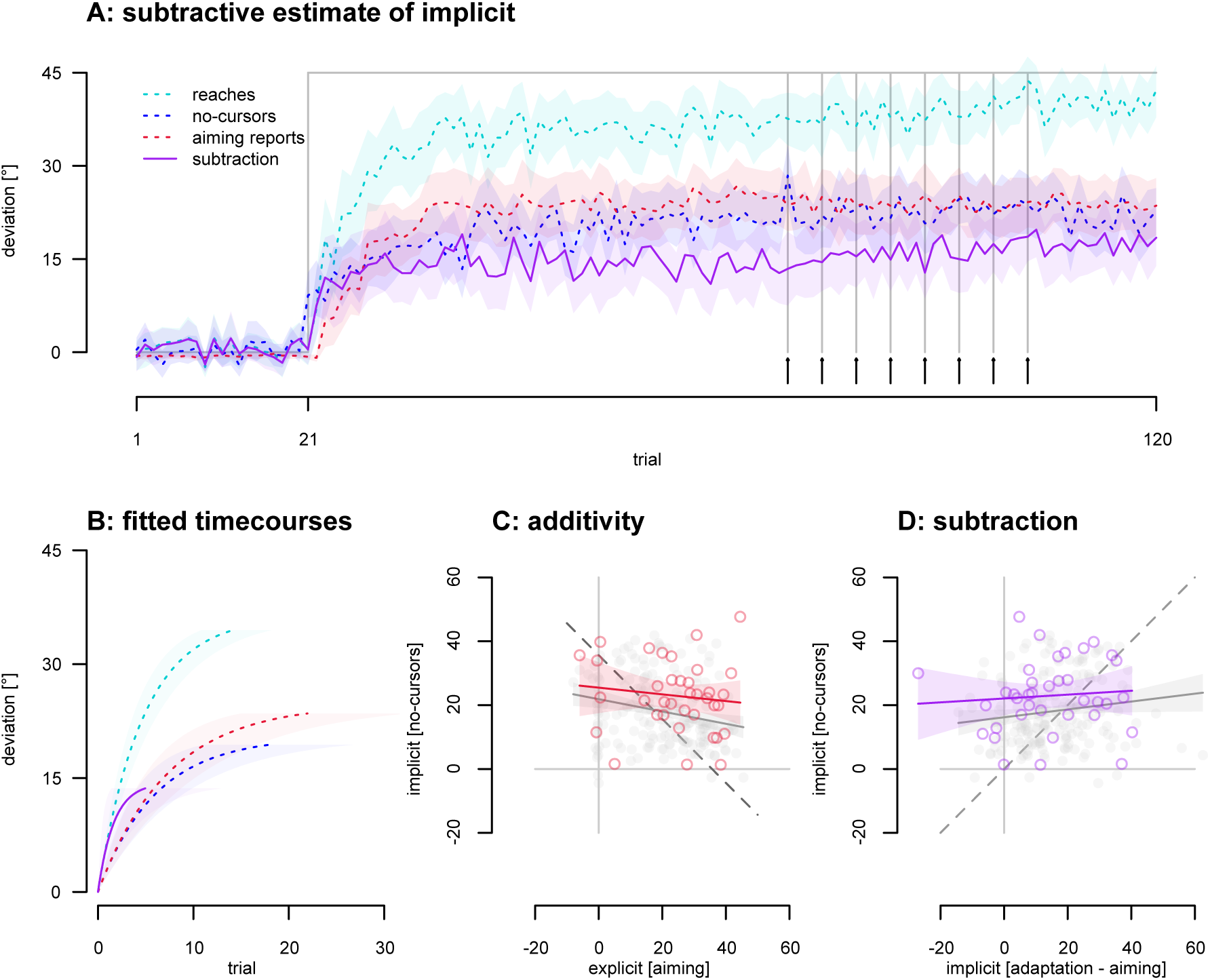
Subtraction comparisons. Shaded regions indicate 95% confidence intervals. **A.** Adaptation across trials, with eight aiming trials in the second half of the aligned phase (indicated by arrows and vertical lines). **B.** Fitted exponential curves. **C.** Individual data scatter plot with regression lines depicting the relationship between implicit and explicit learning processes when using aiming. Each dot represents a participant. Light gray represents data from all other participants doing a 45° rotation. **D.** Individual data scatter plot with regression lines depicting the relationship between implicit and explicit learning processes when using subtraction. Each dot represents a participant. Light gray represents data from all other participants doing a 45° rotation.

**Fig 10.**
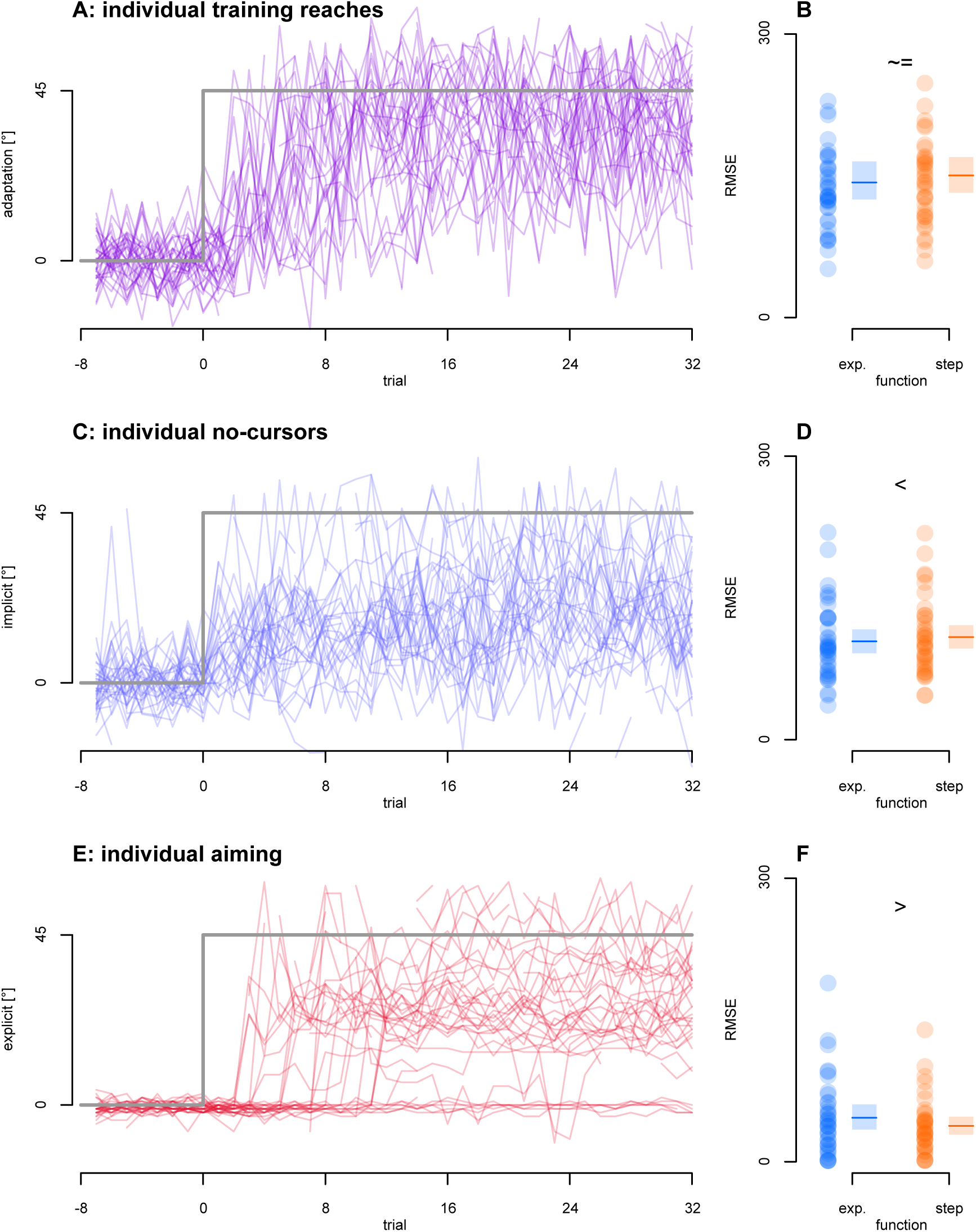
Individual participants’ timecourses. The first 32 trials after the rotation is introduced (trial 0) of overall, implicit and explicit adaptation (on the left), as well as goodness-of-fit estimates (root mean square errors, RMSE) of exponential- and step functions fitted to these timecourses (on the right). **A., B.** Overall adaptation. **C., D.** Implicit adaptation. **E., F.** Explicit adaptation.

**Fig 11.**
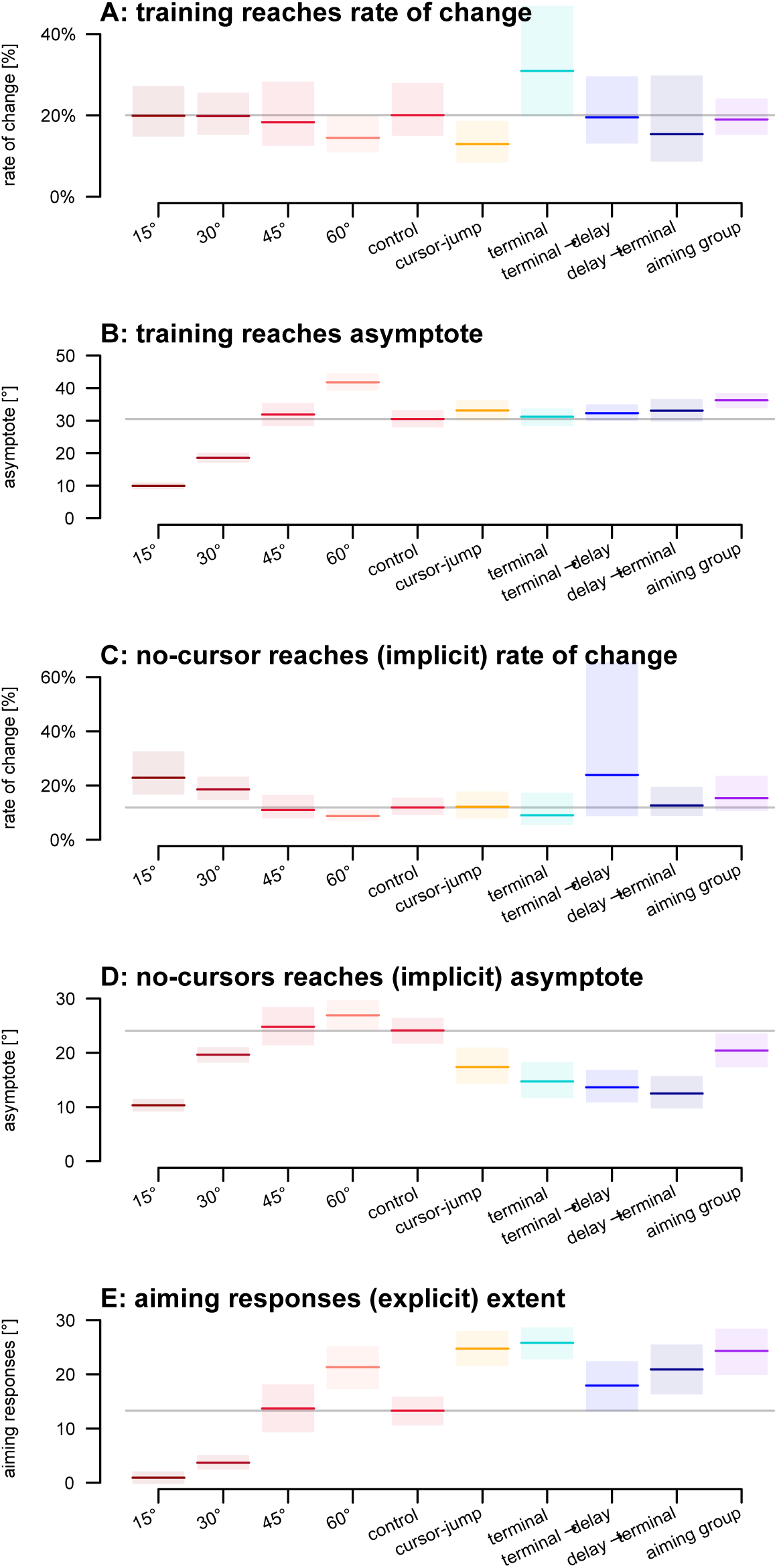
Summary figure of all groups and adaptation indicators. Colored lines indicate group averages and shaded areas denote bootstrapped 95% confidence intervals of the mean. The gray lines show the averages of the control group for comparison. **A.** Reach training rates of change. **B.** Reach training asymptotes. **C.** Implicit no-cursor rates of change. **D.** Implicit no-cursor asymptotes. **E.** Explicit re-aiming extents.

## Discussion

Our study sought to investigate the time-course of implicit adaptation during classical, or movement-contingent, visuomotor adaptation using interleaved no-cursor trials to gauge if implicit adaptation is 1) a slow process in adaptation, 2) affected by rotation size, 3) modulated by conditions that (mostly) increase explicit adaptation, and 4) linearly additive with explicit adaptation. Using no-cursor reach aftereffects after every reach training trial, we can map out the speed of implicit adaptation with high temporal precision. All the non-control conditions in experiments 2 through 4 involve conditions that usually evoke more explicit strategies, which could then decrease implicit adaptation – or slow it down. Our results challenge the traditional notion that implicit adaptation is a slow process, as we found that implicit learning processes emerge much faster than previously assumed in all conditions (Fig 11) and about as fast as explicit strategies. Using interleaved no-cursor trials, we validate the efficacy of this method through the expected effects of increasing rotation size. We observed that cursor-jump feedback and terminal feedback primarily enhance explicit adaptation while having minimal impact on the speed and asymptote of implicit adaptation. For the group that did continuous aiming reports, we find no discernible effect on implicit adaptation, highlighting a parallel development of implicit and explicit adaptation at comparable speeds. The rapid emergence of aftereffects we found was robust to our various feedback types and the aftereffects of implicit learning were observed to develop within the first few training trials, and reaching asymptote within 20 training trials for all conditions and as few as 10 trials for half of our conditions. This indicates that unconscious adaptation can occur very rapidly.

Our experimental approach represents an advancement in the study of implicit learning during *classical* or movement-contingent adaptation by introducing key improvements in measurement and analysis. Like in our previous three papers [4,24,25], we consistently measure aftereffects to capture the residual deviations in reaching movements even after the feedback is removed or returned to normal, providing a more comprehensive approach that ensures a robust assessment of the natural time course of implicit adaptation. Traditional measures of implicit adaptation consist of reach aftereffects after many trials, masking initial changes. More recent measures either rely on subtraction explicit adaptation from total adaptation, which we have shown not to be ideal [5], or on clamped error feedback combined with instructions to ignore this feedback, probably suppressing learning. Instead we used an independent and direct measurement. By avoiding the subtraction method, which assumes additivity between the two processes, we can better assess each process’ individual contributions as well as potential interactions. The limitation of employing a subtractive approach is further demonstrated in Fig 9F, where it is evident that aiming responses at the individual level do not exhibit exponential behavior. Instead, individuals seem to become cognizant of the perturbation at various discrete points during the learning process. This is in contrast to the gradual increments typically observed in the implicit processes of individuals with no strategy, as seen in the interleaved reach aftereffects in Fig 9E, where the expected progression of an unconscious process is evident.

The inclusion of no-cursor trials may have introduced some interference with overall adaptation. In a previous study, we found that interleaving no-cursor trials led to a lower asymptote at 77%, compared to 96% when there were no interleaved no-cursor reaches but just a gap in time [4]. This suggests that interference from no-cursor trials had some effect on overall adaptation. However, in this previous study we observed higher rates of change, and that study used passive movements for the return to home in all trial types, whereas here we used active movements. Since the previous study showed that active interlaced movements reduced learning somewhat, this may explain some of the differences in findings between our two studies. More precisely: the inclusion of the no-cursor trials to measure implicit adaptation probably slowed down implicit adaptation and overall adaptation. Importantly, we did not include any groups without no-cursors, so that the effect of the interlaced no-cursor trials was present in all groups, and implicit adaptation was still quite fast in all groups. This likely slowdown of implicit adaptation in the current study should be taken into account when making comparisons with other research.

Finally, it is possible that participants did not fully disengage their strategy when switching from cursor-to-target training trials to no-cursor hand-to-target trials. If so, larger strategies should lead to larger no-cursor deviations. The Rotation Size experiment shows an effect like this. Though increases in implicit adaptation there could also just be driven by the increasing rotations, rather than by carry-over effects. The other groups, however, all used the same rotation size, so carry-over of greater aiming should have produced larger no-cursor deviations. Instead these were smaller. This suggests that our measure of implicit adaptation is only minimally affected.

In our study, we also investigated the relationship between rotation size and the time-course of implicit learning. Not surprisingly, larger rotations led to a proportionally larger overall adaptation extent. However, in contradiction to earlier work [6] including our own [7], we found that the extent of reach aftereffects did vary a bit with rotation size during training, but the difference was only 7° for rotations between 30° and 60° rotations, as compared to 23° of difference for total adaptation. The rate of change or absolute time-course in degrees, both during rotated-reach training, and during subsequent washout with a veridical cursor, did not clearly vary with the size of rotations between 30° and 60°, as illustrated in Fig 3B and D. Only training with a small rotation like 15° led to any differences, which is in line with the idea of capped implicit adaptation.

Comparing the time-course of aftereffects *during* training for different types of visual feedback offers valuable insights into the factors influencing the progression of implicit learning. Our findings support and extend the work of Ruttle et al. (2021) who also examined aftereffects throughout early reach training. Our study demonstrated continuous reach aftereffects at a rate of change of 20.1% (CI 15.0%-28.0%) in contrast to the 56.9% (CI 27.4%–58.5%) we reported earlier [4]. Despite this difference, both studies challenge the traditional notion of slow implicit adaptation, indicating that it *still* occurs at a notably faster pace.

Furthermore, we delved into the influence of feedback type, revealing that terminal and cursor jump feedback both led to smaller implicit reach aftereffects than continuous feedback, suggesting potential competition between explicit strategy engagement and implicit adaptation [26]. Like others, we found that terminal feedback lowered the extent of implicit adaptation, perhaps due to its limited visual cues [8–12]. However, our observations highlight that implicit adaptation can still rapidly emerge within this context. While we did not find a slower rate for cursor jump, we did replicate our earlier finding of reduced aftereffects [15].

Our results indicate that delaying terminal visual feedback does not substantially alter either the magnitude of implicit adaptation (about 14°) or the rate at which it emerges, suggesting that delaying feedback does not diminish implicit adaptation at all. Our finding appears to contradict prior reports that delayed feedback reduces or abolishes implicit adaptation while increasing reliance on explicit strategies [9,11,27,28]. However, closer inspection of these studies reveals that implicit learning was attenuated rather than eliminated. For example, Schween and Hegele (2017) reported aftereffects of 11.1° under continuous feedback, which were reduced to 7.2° and 4.0° with 0.2-s and 1.5-s delays, respectively.

Brudner et al. (2016) likewise found robust immediate aftereffects (∼10°) under both continuous and delayed terminal feedback, with faster washout in the delayed condition. McDougle and Taylor (2019) observed smaller but significant aftereffects (∼2.7°) following 2-s delays, which could have underestimated adaptation due to averaging across many washout trials. Even Tsay et al. (2023), despite concluding that delays of 800 ms abolish implicit learning, present data in which aftereffects in healthy controls—and their confidence intervals—remain clearly above zero. In our study, we directly measured implicit aftereffects during movement-contingent visuomotor adaptation and found that they saturated within ∼20 trials. While their magnitude was reduced relative to continuous feedback, they were unaffected by whether terminal feedback was presented immediately after movement offset or 1.2 s later. Regardless, terminal (with and without a delay) and cursor jump feedback do not influence the rate at which implicit adaptation unfolds.

While our study primarily focused on implicit components of adaptation, we also examined the extent of explicit adaptation. We used an aiming task to determine the explicit contribution to adaptation across our feedback types, and our results suggest that error feedback type (terminal & cursor jump) can increase the amount of explicit control over the task, aligning with previous research [12,15]. Taken together with our measure of aftereffects, we now had direct measurements of both implicit and explicit processes. After performing linear regressions on this data we consistently found a non-additive relationship between implicit and explicit, in line with recent work from our lab [5]. Expanding on aiming, our experiment also explored how taking frequent explicit measurements can affect the progression of implicit learning. Exploring this continuous aiming, we find that explicit aiming judgments do not impact the rate of implicit learning, while it slightly increases the overall extent of adaptation without affecting its speed. Likewise, when plotting at the individual level, it seems that aiming or cognitive strategies tend to manifest discretely, with participants suddenly becoming aware or reporting strategies at some point during early learning. Consequently, when these measures are averaged across participants, they may appear deceptively exponential. Thus, this cognitive component does not appear to resemble an exponential the way that overall adaptation and implicit adaptation does. Combined with our finding that the extent of implicit and explicit adaptation are not additive, this suggests that both implicit and explicit adaptation can develop quite rapidly and to some degree independently of each other, but see [26]. The speed of explicit processes has been extensively explored in the field and is generally agreed to be remarkably fast [12,29–31]. Consequently, our study highlights the importance of considering the speed of implicit adaptation, and that further exploration of implicit learning is warranted.

Additionally, we see in all four experiments that the distribution of the amount of implicit adaptation seems predominantly uni-modal, whereas the level of explicit adaptation may follow a multi-modal distribution. A portion of participants seems not to develop any strategy, whereas others have strategies that fall in clusters. We observed something related previously [5] so this is not wholly unexpected. In this data set we also find that participants appear to develop their explicit strategies in a sudden moment of insight (Fig 10), which could be more characteristic of a cognitive process [32], while the time course of implicit reach aftereffects aligns more with a gradual process. However, instead of speculating on it, we will leave this phenomenon for future investigation.

In conclusion, our study challenges conventional assumptions about the time-course of implicit adaptation during visuomotor tasks. We provide evidence that implicit learning can occur rapidly within the initial stages of training, across different feedback conditions, rotation sizes, and feedback delay timings. We also find that the speed of implicit adaptation was indistinguishable from the speed of explicit adaptation. This has important implications for our understanding of how motor learning processes unfold and interact, and the complex synergy between implicit and explicit components of adaptation. Further research in this direction could offer insights into optimizing motor learning interventions and training strategies.

## Data Availability

Data and analyses are available on Open Science Framework (https://osf.io/ajwyr/).

## Acknowledgements

This work was supported by NSERC for SD; and NSERC for DYPH. The funders had no role in study design, data collection and analysis, decision to publish, or preparation of the manuscript.

## Contributions

BMtH and DYPH. designed the research. SD collected the data. SD, JER and BMtH analyzed the data. SD wrote the manuscript, SD, BMtH and DYPH edited and revised the manuscript. All authors reviewed and approved the manuscript.

